# Macrophage Depletion Protects Against Cisplatin-Induced Ototoxicity and Nephrotoxicity

**DOI:** 10.1101/2023.11.16.567274

**Authors:** Cathy Yea Won Sung, Naoki Hayase, Peter S.T. Yuen, John Lee, Katharine Fernandez, Xuzhen Hu, Hui Cheng, Robert A. Star, Mark E. Warchol, Lisa L. Cunningham

**Affiliations:** Laboratory of Hearing Biology and Therapeutics, National Institute on Deafness and Other Communication Disorders (NIDCD), NIH, Bethesda, Maryland, USA; Renal Diagnostics and Therapeutics Unit, National Institute of Diabetes and Digestive and Kidney Diseases (NIDDK), NIH, Bethesda, Maryland, USA; Washington University, Department of Otolaryngology, School of Medicine, Saint Louis, MO; Bioinformatics and Biostatistics Collaboration Core, National Institute on Deafness and Other Communication Disorders (NIDCD), NIH, Bethesda, Maryland, USA

**Keywords:** Cisplatin, Hair cells, Ototoxicity, Nephrotoxicity, Auditory Brainstem Response (ABR), Macrophages, Ablation, CSF1R, PLX3397, Kidney

## Abstract

Cisplatin is a widely used and highly effective anti-cancer drug with significant side effects including ototoxicity and nephrotoxicity. Macrophages, the major resident immune cells in the cochlea and kidney, are important drivers of both inflammatory and tissue repair responses. To investigate the roles of macrophages in cisplatin-induced ototoxicity and nephrotoxicity, we used PLX3397, an FDA-approved inhibitor of the colony-stimulating factor 1 receptor (CSF1R), to eliminate tissue-resident macrophages during the course of cisplatin administration. Mice treated with cisplatin alone (cisplatin/vehicle) had significant hearing loss (ototoxicity) as well as kidney injury (nephrotoxicity). Macrophage ablation using PLX3397 resulted in significantly reduced hearing loss measured by auditory brainstem responses (ABR) and distortion-product otoacoustic emissions (DPOAE). Sensory hair cells in the cochlea were protected against cisplatin-induced death in mice treated with PLX3397. Macrophage ablation also protected against cisplatin-induced nephrotoxicity, as evidenced by markedly reduced tubular injury and fibrosis as well as reduced plasma blood urea nitrogen (BUN) and neutrophil gelatinase-associated lipocalin (NGAL) levels. Mechanistically, our data suggest that the protective effect of macrophage ablation against cisplatin-induced ototoxicity and nephrotoxicity is mediated by reduced platinum accumulation in both the inner ear and the kidney. Together our data indicate that ablation of tissue-resident macrophages represents a novel strategy for mitigating cisplatin-induced ototoxicity and nephrotoxicity.

**Brief summary:** Macrophage ablation using PLX3397 was protective against cisplatin-induced ototoxicity and nephrotoxicity by limiting platinum accumulation in the inner ear and kidney.

## Introduction

Cisplatin is a platinum-based chemotherapeutic drug that is widely used and highly effective in treating a variety of solid tumors, including testicular, bladder, lung, stomach, head and neck, and ovarian cancers, in both pediatric and adult patients (*1*). It exerts its anti-tumor activity by forming DNA cross-links that interfere with replication, transcription, and DNA repair mechanisms, causing DNA damage and subsequent apoptosis in cancer cells (*1, 2*). Cisplatin therapy is associated with significant side effects, including ototoxicity, nephrotoxicity, and myelosuppression, which can be dose-limiting and can compromise therapeutic outcomes as well as quality of life for cancer survivors (*3–6*).

Cisplatin is the most ototoxic drug in clinical use, resulting in bilateral, progressive, and permanent sensorineural hearing loss in up to 60% of treated adults (*7–9*) and 70% of treated children (*10–13*). Cisplatin results in the death of mechanosensory hair cells in the inner ear that are required for hearing function (*14*). Recently, the FDA approved the use of sodium thiosulfate (STS) as an otoprotective agent for mitigating cisplatin-induced ototoxicity in pediatric patients with localized, non-metastatic solid tumors (*15–18*). Despite the significant progress in ototoxicity research, there are currently no effective treatment strategies for cisplatin-induced ototoxicity in adults or in children with metastatic cancers. In addition to ototoxicity, cisplatin also results in kidney damage and renal dysfunction. Approximately 30-40% of cancer patients treated with cisplatin develop acute kidney injury (*19–21*). In addition, patients without acute kidney injury remain susceptible to the development of chronic kidney disease both during and after the discontinuation of cisplatin treatment. Currently, although hydration can rehabilitate acute renal damage, there are no effective pharmacological treatments available for cancer patients receiving cisplatin-based therapies who develop chronic kidney disease without acute kidney injury (*22–24*). Thus, there is an unmet clinical need for novel treatments that limit or prevent cisplatin-induced ototoxicity and nephrotoxicity.

Cochlear macrophages are the major innate immune cells in the cochlea and act as important drivers of both inflammatory (*25*) and tissue repair responses (*26, 27*) after cochlear injury. Macrophages account for >95% of CD45+ leukocytes in the adult mouse cochlea (*28*) and are also found in large numbers in the human cochlea (*29–31*). Under steady-state conditions, macrophages are present in the osseous spiral lamina, basilar membrane, spiral ganglion neurons (SGNs), spiral ligament, and stria vascularis (*28, 32, 33*), while the organ of Corti is mostly devoid of macrophages (Figure 1C”) (*28*). The stria vascularis contains the blood supply of the cochlea as well as a unique subset of tissue-resident macrophages, the perivascular macrophages (PVMs), that are closely associated with the blood vessels and function as part of the blood-labyrinth barrier (BLB) (*34–36*). The BLB between the lumen of the vasculature and the inner ear fluid spaces tightly controls the exchange of substances circulating in the bloodstream and those in the inner ear (*37*). PVMs are important for BLB function and regulate the BLB permeability through either direct physical interaction with the capillaries or by secreting soluble factors that act on vascular endothelial cells that line the capillary lumen (*34, 38, 39*). Insults to the cochlea, such as exposure to loud noise (*40–43*), bacterial or viral infections (*44–46*), or insertion of a cochlear implant (*47–49*), can activate tissue-resident macrophages, which can upregulate inflammatory cytokines, phagocytose tissue debris, or induce infiltration of peripheral immune cells to the site of injury (*50*). However, the roles of cochlear macrophages in cisplatin-induced ototoxicity are unknown.

**Figure 1.**
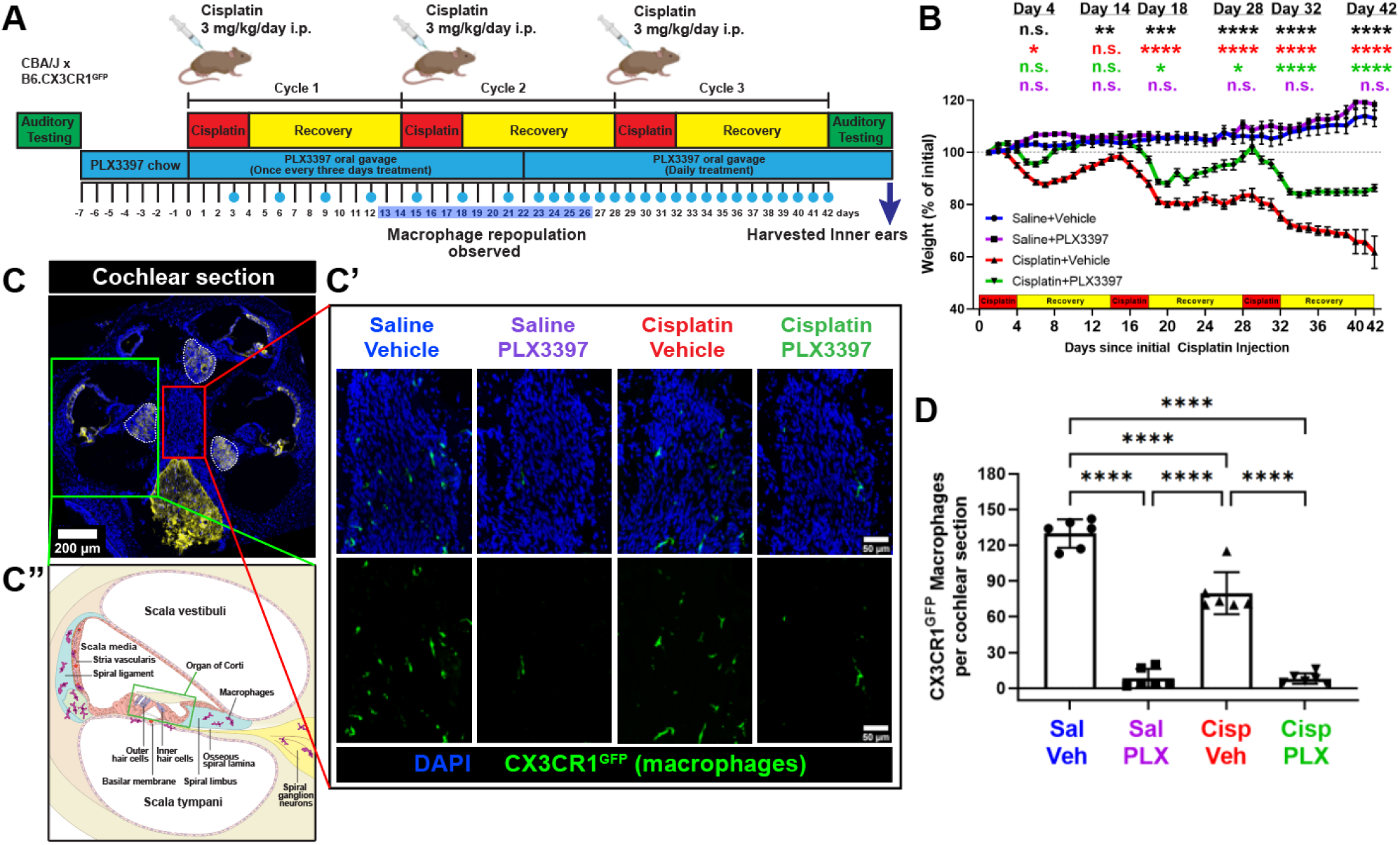
Initial protocol in Experiment 1 resulted in macrophage ablation via PLX3397 with partial repopulation. (A) Experimental design showing auditory tests, PLX3397 treatment, and three cycles of cisplatin administration. Mice received PLX3397-formulated chow for seven days to facilitate macrophage depletion. During cisplatin administration, mice received PLX3397 via oral gavage once every three days. Upon observation of macrophage repopulation on day 14 (see Supplemental Figure 1), daily treatment of PLX3397 via oral gavage was initiated. The days on which mice received PLX3397 via oral gavage are denoted with blue circles. (B) Mice treated with the combination of cisplatin and PLX3397 exhibited significantly less weight loss compared to mice treated with cisplatin and vehicle. *Mean±SEM, n=10-16 mice per experimental group*. *Statistical comparisons (asterisks or n.s.) are color-coded as described in Methods section*. (C) Cochleae harvested after endpoint auditory tests were immunolabeled for Kir4.1 (yellow) to visualize cochlear structures and GFP to visualize CX3CR1^GFP^-positive macrophages (green). (C’) Representative confocal images (cochlear modiolus; from red box in panel C) and (D) quantification of macrophages per cochlear section. PLX3397 resulted in >93% depletion of all macrophages. (C’’) Schematic diagram of macrophage distribution in the middle cochlear turn (from green box in panel C). *Scale bar, 200 μm (cochlear section) and 50 μm (mid-modiolar section).* (D) Quantification of macrophages per cochlear section. *Mean±SD, n=6 cochleae per experimental group. One-way ANOVA with Tukey’s multiple comparisons test*.

Kidney resident macrophages are critical components of the innate immune system in the kidney, and they are involved in maintaining tissue homeostasis, immune surveillance, response to tissue injury, and wound healing (*51, 52*). Kidney resident macrophages account for approximately 50-60% of CD45+ leukocytes in the mouse kidney (*53*) and include heterogenous populations with varying phenotypes and functions depending on their location within the kidney (*54, 55*). Kidney resident macrophages are found closely associated with capillaries that surround the tubules and glomeruli (*53*) and are involved in the regulation of tubular epithelial cell function or induction of apoptosis during tubulointerstitial inflammation (*56–58*). Moreover, emerging evidence suggests that kidney resident macrophages are involved in mediating cisplatin-induced renal injury (*59*). However, the precise mechanisms by which renal macrophages contribute to cisplatin-induced nephrotoxicity are not fully understood.

The goals of the present study were to examine the roles macrophages play in cisplatin-induced ototoxicity and nephrotoxicity and to identify the mechanism(s) underlying their activity in response to cisplatin using a mouse model of cisplatin ototoxicity that results in hearing loss that is similar to that observed clinically (*60*). We used a pharmacological inhibitor of colony-stimulating factor-1 receptor (CSF1R) to achieve prolonged depletion of tissue-resident macrophages *in vivo*. Colony-stimulating factor-1 (CSF1) signaling through its receptor CSF1R promotes proliferation, differentiation, and survival of tissue-resident macrophages (*61–64*). PLX3397 is a small-molecule inhibitor of CSF1R that penetrates the blood-brain barrier (BBB) and results in the elimination of 95-99% of all microglia in the CNS (*61, 63–66*). PLX3397-induced macrophage/microglia ablation in the CNS is achieved while minimally affecting circulating monocytes or other immune cells (*62, 67, 68*). We administered PLX3397 seven days prior to and throughout the cisplatin administration period to examine the roles of macrophages during cisplatin treatment. Our data indicated that >88% of tissue-resident macrophages were eliminated in both the cochlea and kidney after PLX3397 treatment. Macrophage ablation resulted in significant protection against cisplatin-induced hearing loss and kidney injury, in part by limiting cisplatin accumulation in the cochlea and the kidney. Our findings strongly suggest that tissue-resident macrophages contribute to cisplatin entry and accumulation into the cochlea and the kidney. We propose macrophage-based immunotherapy as a novel approach to prevent cisplatin ototoxicity and nephrotoxicity in cancer patients undergoing cisplatin therapy.

## Methods

### Animals

Studies were performed using adult mice of F1 offspring from CX3CR1^GFP/GFP^ (C57BL/6 background, The Jackson Laboratory, stock #005582) cross bred with CBA/J (The Jackson Laboratory, stock #000656). C57BL/6 WT mice develop sensorineural hearing loss as early as 3-4 months of age, much earlier than other mouse strains such as CBA/J (*69, 70*). Therefore, this mouse breeding strategy was employed to overcome the early onset hearing loss inherent in the C57BL/6 WT mouse strain while maintaining Enhanced Green Fluorescent Protein (EGFP) expression in macrophages. CX3CR1^GFP/GFP^ (C57BL/6 background) homozygous mice express EGFP under the control of the endogenous Cx3cr1 locus, where the Cx3cr1 gene is replaced with EGFP resulting in mice that function as CX3CR1 knockout mice (*71*). Here, we utilized a CX3CR1^GFP/+^ heterozygous mice in which one copy of the Cx3cr1 gene is replaced with EGFP while the other copy is still functional. This allows us to maintain the function of CX3CR1 in these mice while also visualizing CX3CR1-expressing cells through the expression of EGFP. Heterozygous mice were genotyped by PCR amplification of genomic DNA extracted from tail snips as recommended by The Jackson Laboratory (https://www.jax.org/jax-mice-and-services/customer-support/technical-support/genotyping-resources/genotyping-protocol-database). The F1 mouse strain with EGFP-labeled macrophages is referred to as CBAJ/CX3CR1^GFP/+^.

### Study design

The study was performed under NIH NIDCD/NINDS Animal Care and Use Committee-approved protocol (#1327). Male and female CBAJ/CX3CR1^GFP/+^ mice (9-14 weeks old) were housed individually and assigned to one of four treatment groups: 1) Saline/Vehicle, 2) Saline/PLX3397, 3) Cisplatin/Vehicle, or 4) Cisplatin/PLX3397. Mice were given *ad libitum* access to food and water. Cisplatin-treated mice received three cycles of once-daily cisplatin (3 mg/kg i.p.) for 4 days followed by a 10-day recovery period (3mg/kg x 4 days x 3 cycles = 36 mg/kg cumulative dose) as described previously (*60*). Control animals received same volume of sterile 0.9% saline. Cisplatin-treated mice received supportive care: (1) daily 1 ml of saline subcutaneous injections (s.c.) in the morning and normasol (Hospira, Inc., San Clemente, CA, USA) in the afternoon; (2) 0.3ml STAT® high-calorie liquid supplement (PRN Pharmacal, Pensacola, FL, USA) twice daily to maintain body weight. Mice were monitored twice daily for changes in overall health or activity using a daily body condition score, for overall condition (muscular tone, body fat content, coat maintenance, and overall energy level) (*72*).

To achieve experimental macrophage depletion, PLX3397 was administered via a chow formulation followed by oral gavage. To prepare PLX3397 chow, PLX3397 was purchased from MedChemExpress Inc. (HY-16749, Monmouth Junction, NJ, USA) and sent to Research Diets Inc. (New Brunswick, NJ, USA) to formulate in AIN-76A standard chow at 660ppm (660mg/kg). On average, mice consume approximately 4g of chow per day, resulting in a daily intake of 2.64mg of PLX3397. Standard AIN-76A rodent chow without PLX3397 (D10001, Research Diets Inc.) was purchased to serve as a control chow.

To prepare 50mg/kg PLX3397 solution for oral gavage, PLX3397 was dissolved in 100% PEG300 (202371, Sigma-Aldrich) to make a concentration of 25mg/ml and incubated on a rotating platform at room temperature for 15-30 minutes. Subsequently, this mixture was subjected to sonication in an ultrasonic bath (Branson 1510) for 20-40 minutes to ensure complete solubilization of PLX3397 and stored at −20°C until use. Separately, 10% Tween 80 stock was prepared in saline (0.9% NaCl) and stored at room temperature until use. On the day of treatment, 25mg/ml PLX3397/PEG300 solution was mixed with 10% Tween 80/Saline to make a final concentration of 10mg/ml PLX3397 in 40% PEG300, 4% Tween 80, and 56% saline diluent. Animals assigned to vehicle treatment were given comparable volumes of the diluent without PLX3397. Although dimethyl sulfoxide (DMSO) is commonly used as a component of the solvent to dissolve PLX3397 for oral gavage (*67, 73–76*), it was excluded from our study due to its known ototoxic properties (*77, 78*). Mice receiving a dose of 50mg/kg PLX3397 via oral gavage received an estimated dose of 1.25mg PLX3397 per oral gavage for an average mouse weighing 25g, which is equivalent to a concentration of approximately 310mg/kg of PLX3397 in the formulated chow.

Two independent experiments were performed. In both experiments, auditory tests were performed prior to the start (baseline) and at the end (endpoint) of the cisplatin administration protocol. Following baseline auditory tests, chow containing 660ppm PLX3397 (or control AIN-76A rodent chow) replaced the regular mouse diet for seven days to ablate (or maintain) macrophages. Following the 7-day pretreatment, mice were returned to the regular rodent chow (5L8F) and underwent the cisplatin administration protocol as they continued to receive PLX3397 via oral gavage. Mice receiving cisplatin demonstrate reduced appetite; therefore, PLX3397 chow feeding could have resulted in insufficient drug administration. Thus, vehicle or 50mg/kg PLX3397 was administered via oral gavage during the cisplatin administration protocol to ensure accurate drug administration. In Experiment 1, vehicle or 50mg/kg PLX3397 was administered by oral gavage once every three days until day 22 of the 42-day cisplatin administration protocol. On this day we observed repopulation of the cochlear macrophages; therefore, PLX3397 administration frequency was increased to daily (Figure 1A, Supplemental Figure 1). Since we had observed some macrophage repopulation in Experiment 1, we conducted a second study (Experiment 2), in which vehicle or 50mg/kg PLX3397 was administered daily throughout the cisplatin administration protocol to ensure that macrophages remained ablated throughout the experiment (Figure 4). In each experiment, endpoint auditory tests were conducted at the end of the 42-day cisplatin administration protocol, and PLX3397 administration via oral gavage was continued during the auditory testing phase until mice were euthanized for tissue harvest.

### Auditory testing

Prior to the start of PLX3397 chow pre-treatment and at the end of the cisplatin administration protocol, mice underwent two types of auditory tests: auditory brainstem responses (ABRs; measure hearing sensitivity) and distortion product otoacoustic emissions (DPOAEs; indirectly measure outer hair cell function).

Mice were anesthetized with ketamine (Putney Inc., Portland, ME; 100mg/kg i.p.) and xylazine (Akorn Inc., Lake Forest, IL; 10mg/kg i.p.). Additional injections at 1/3-1/2 of the original dose were administered if needed. Animals were placed on a temperature-controlled (37°C) heating pad (World Precision Instruments T-2000, Sarasota, FL, USA) during auditory testing inside a noise-cancelling chamber (Acoustic Systems, Austin, TX, USA). ABRs and DPOAEs were recorded from the left ear of each mouse using Tucker Davis Technologies (TDT) hardware (RZ6 Processor) and software (BioSigRZ).

For ABR measurements, subcutaneous needle electrodes (Rhythmlink, Columbia, SC, USA) were placed behind left pinna of the test ear (reference), vertex (active), and near the tail of the mouse (ground). Tone-burst stimuli (Cos2, 3 msec, 0.5 msec rise/fall in alternating polarity) were presented at a rate of 29.9/sec at 8, 11.2, 16, 22.4, 32, and 40 kHz starting at 90dB SPL. At each sound level, 1024 waveforms were averaged, amplified (20x), and filtered (HP: 300Hz, LP: 3kHz, NT: 60Hz). At near-threshold sound pressure levels (SPL), the ABR waveforms were recorded twice, and two waveforms were superimposed for comparison. ABR threshold was defined as the lowest stimulus intensity that resulted in a reproducible waveform displaying identifiable peaks.

DPOAEs were measured in response to two primary pure tones, f1 and f2, generated by Multi Field 1 speakers. Two primary tones were presented at 6 frequency pairs, where f2 corresponded to ABR test frequencies (f2 = 8, 11.2, 16, 22.4, 32, 40kHz; f2/f1 = 1.25). The sound level was increased in 5dB steps from 30dB to 90dB. At each sound level, 512 responses were averaged. DPOAE at 2f1-f2 was recorded in the mouse inner ear canal using an ER-10B+ microphone (Etymotic, Elk Grove Village, IL, USA) connected to a modified pipette tip to fit the mouse external ear canal. Biological noise floors and amplitudes were calculated for each treatment group and plotted relative to each other.

### Blood and Tissue collection

All experimental animals were anesthetized by injecting a cocktail of ketamine (120mg/kg) and xylazine (25mg/kg). After achieving surgical level of anesthesia (verified by the lack of hind paw pinch reflex and eyelid reflex), an incision was made along the abdomen. The abdominal cavity was exposed and the widest part of the inferior vena cava between the kidneys was located. A heparin-coated 28G needle (0.2% Heparin solution, 07980, STEMCELL Technology Inc.) in a 1mL syringe was carefully inserted into the vein to withdraw blood. Blood was transferred into an Eppendorf tube on ice, spun at 3000rpm for 25 minutes at 4°C using a refrigerated benchtop centrifuge (Eppendorf centrifuge 5427R), and stored at −80°C until further use.

Following blood collection, cardiac perfusion with 1X PBS (diluted in mqH_2_O from 10X PBS, 70011-044, Thermo Fisher) was performed. The thoracic cavity was opened by incising through the rib cage and diaphragm, and the heart was exposed. The right atrium was opened by incision; a 23-25G butterfly needle was inserted into the left ventricle, and the body was perfused with 1X PBS via the blood flow. After cardiac perfusion, mice were decapitated using surgical scissors, and the inner ears, kidneys, and spleen were collected for further analyses. Inner ears were dissected from the temporal bone; a small hole was opened in the apex of the cochlea using a 27G insulin needle, and the cochlea was perfused with 4% PFA (diluted from 16% PFA, 5710-S, Electron Microscopy Sciences) through the round window, oval window, and the apical hole for complete fixation of the tissue. Inner ears were then post-fixed for 5-6 hours at room temperature. Following fixation, inner ears were washed with 1X DPBS (14190-250, Thermo Fisher) and decalcified in 0.5M EDTA solution (pH 8.0) for 26 hours at room temperature. Inner ears were washed again with 1X DPBS and stored at 4°C until dissection. Left kidneys were cut in the horizontal plane into three pieces, frozen immediately in dry ice, and stored at −80°C. Right kidneys were dissected out and renal capsules were removed and fixed in 10% neutral buffered formalin and were prepared for histology. Spleens were collected to examine the impact of PLX3397 on peripheral immune cells using flow cytometry.

### Microdissection, cryosectioning, and staining (inner ear)

For wholemount dissections of the cochlea, the stria vascularis was peeled from the cochlear lateral wall. Organ of Corti and SGNs were cut from the surrounding tissues and microdissected into five pieces (apex, mid-apex, mid-base, base, base-hook) spanning all cochlear regions from base to apex.

For mid-modiolar sections of the inner ear, fixed and decalcified inner ears were cryoprotected in 15% sucrose/DPBS overnight, 20% sucrose/DPBS overnight, and 30% sucrose/DPBS for two days at 4°C. Subsequently, inner ears were immersed in 1:1 mixture of 30% sucrose and super cryoembedding medium (SCEM) (C-EM001, Section-Lab Co. Ltd., Hiroshima, Japan) for ∼1 hour at room temperature. The tissues were then embedded in 100% SCEM within a cryomold biopsy square (NC9806558, Fisher) and snap-frozen in 2-methylbutane/dry ice. A cryostat microtome (CM3050S, Leica, Vienna, Austria) was used to cut 14μm thick mid-modiolar frozen tissue sections. Inner ear sections were dried overnight at room temperature and stored in −80°C until further processing.

For immunostaining, dissected cochlear or stria vascularis wholemounts were incubated in blocking/permeabilization buffer (0.3% Triton X-100, 3% NGS, and 2% BSA in 1X DPBS) for 1-2 hours at room temperature. Frozen inner ear sections were rehydrated in PBS and then incubated in blocking buffer. Tissues were probed with primary antibodies (diluted in blocking buffer) at room temperature for 1 hour or overnight at 4°C. Subsequently, the tissues were incubated in species/isotype-matched secondary antibodies conjugated with Alexa Fluor (Thermo Fisher) or TRITC (Southern Biotech) at room temperature and stained for nuclei using Hoechst 33342 (H3570, Thermo Fisher). Information on the antibodies (target, species, and source) and reagents is in Table 1. For CtBP2 and GluR2 staining, cochlear wholemount tissues were incubated in blocking buffer (1% Triton X-100, 5% NGS, and 1% BSA in 1X DPBS) at room temperature for 1 hour followed by primary antibody incubations in antibody dilution buffer (1% Triton X-100 and 1% NHS, in 1X DPBS) at 37°C overnight in a humidifying container. Following primary antibody staining, cochlear tissues were incubated in secondary antibodies diluted in blocking buffer and incubated at 37°C for 1 hour (*79*). After immunostaining, tissues were mounted with either Vectashield (H-1000, Vector Biolabs) or Prolong Gold (P36934, Thermo Fisher) antifade mounting medium onto glass slides, cover slipped, and sealed with nail polish. The stria vascularis was mounted with the marginal cells oriented towards the cover glass.

### Microscopy and data analyses (Inner ear)

For cochlear wholemounts, low magnification confocal images of microdissected cochlear pieces were taken and imported to ImageJ (Fiji; version 2.1.0; NIH, Bethesda, MD). Cochlear length was measured to subsequently convert cochlear locations to cochlear frequencies using ImageJ/Plugin/Tools/Measure_line.class (downloaded from https://www.masseyeandear.org/research/otolaryngology/eaton-peabody-laboratories/histology-core) (*80, 81*). The cochlear frequency map was used to acquire high-resolution confocal z-stack images (Airyscan) from specific cochlear regions corresponding to frequencies of 8, 11.2, 16, 22.4, 32, 40, 63kHz along the basilar membrane in each cochlea. High-resolution confocal-z-stack images (step size of 0.6μm) of hair cells and synapses were acquired using a 63x (1.4 N.A. Oil DIC M27) Plan-Apochromat oil-immersion objective lens with a 0.9X optical zoom at a resolution of 2048×2048 pixels, with a scanning speed of 2.05μsec/pixel. The imaging was performed on an Axiovert 200M inverted microscope with a confocal scan head (Carl Zeiss Microscopy LSM980) equipped with an Airyscan detection unit, all controlled by Zen Black v2.3 software.

For SGN and macrophage quantification in cochlear sections, images encompassing the entire cross-section of the cochlea were obtained with a Zeiss LSM980 microscope through a series of overlapping images (6 or 8 tiles, 10% overlap). Subsequently, these tiled images were stitched together to generate a single image of the whole cochlear section, through the tiling and stitching function in the Zen Black v2.3 software. Images were acquired using a Plan-Apochromat 20x (0.8 N.A. M27) objective lens at a resolution of 1024×1024, with a scanning speed of 0.51μsec/pixel. The stria vascularis wholemounts were imaged using a 20x objective lens (0.8 N.A.) with a 1.1x optical zoom at a resolution of 2048×2048 pixels, with a scanning speed of 2.05μsec/pixel (Airyscan super-resolution).

Outer hair cells (OHCs), inner hair cells (IHCs), spiral ganglion neurons (SGNs), and macrophages were manually counted using the ImageJ software (ImageJ/Point tool) (NIH, Bethesda, MD, USA). IHC synapses were semi-automatically quantified using Imaris x64 9.2.1 and Imaris File Converter x64 9.1.2 (Oxford Instruments, Abingdon, Oxfordshire, England). OHCs, IHCs, and IHC synapses were quantified within the span of 147.90μm. The total numbers of synaptic puncta (CtBP2 and GluR2 staining) were divided by the number of IHCs, to gain the number of puncta associated with each IHC. SGNs in Rosenthal’s canal were quantified from mid-modiolar sections of the inner ear and total number of SGNs were normalized to the area of Rosenthal’s canal to obtain density measurements (SGNs per 10,000μm^2^). The SGN density from each of the three regions (apex, mid, and base) of the cochlea were averaged separately. Macrophages in the osseous spiral lamina of cochlear wholemounts were quantified using a region of interest (ROI) measuring 180μmx180μm ROI. Perivascular macrophages (PVMs) in the stria vascularis wholemounts were quantified in the entire image frame (20X objective zoom, 1.1X digital zoom, Airyscan, 2024×2024 pixels).

### Morphological evaluation (kidney)

After harvesting under anesthesia, kidney specimens were fixed with 10% formalin and subsequently embedded in paraffin. To assess tubular injury, sections (4µm thickness) were stained with Periodic Acid-Schiff (PAS), while Masson-Trichrome (MT) reagent was used to evaluate renal fibrosis. Tubular injury and interstitial fibrosis were evaluated by a blinded observer in 10 randomly selected non-overlapping fields at 400x magnification in each section of cortico-medullary junction. This region contains proximal tubule S3 segments, which are the most vulnerable to cisplatin-induced injury (*82*). As previously described (*83*), tubular injury judged by tubular atrophy, tubular dilation, protein casts, tubular necrosis, and brush border loss was rated on the following scoring system: 0, 0%; 1, 1–25%; 2, 26–50%; 3, 51–75%; 4, 76–100%. The fibrotic area was calculated using Fiji/ ImageJ software (NIH, Bethesda, MD, USA).

### Immunohistochemistry (kidney)

Formalin-fixed paraffin embedded kidney tissues were sectioned, deparaffinized, and subjected to antigen retrieval in sodium citrate buffer (10mM sodium citrate, 0.05% Tween 20, pH 6.0) for 20 minutes at 98°C using a microwave. Endogenous peroxidase activity was blocked with 0.3% hydrogen peroxide in methyl alcohol for 15 minutes. After blocking with goat serum, the specimens were incubated overnight at 4 °C with 5µg/mL rabbit anti-GFP antibody (ab290, Abcam, Cambridge, MA, USA). Subsequently, horseradish peroxidase-conjugated goat anti-rabbit antibody (Agilent Dako, Santa Clara, CA, USA) was applied to sections and incubated for 1 hour at room temperature. Sections were developed using 3,3′-diaminobenzidine tetrahydrochloride (Sigma-Aldrich, St. Louis, MO, USA), and then counterstained with hematoxylin. Positively stained cells were counted in 10 randomly selected non-overlapping fields at 400x magnification of cortico-medullary junction.

### Measurement of kidney function and injury marker

Blood urea nitrogen (BUN) was measured from 2µl plasma using a Quantichrom^TM^ Urea Assay Kit (Bioassay system, Hayward, CA, USA). Plasma neutrophil gelatinase-associated lipocalin (NGAL) was assessed with enzyme-linked immunosorbent assay (R&D Systems, Minneapolis, MN, USA) according to the manufacturer’s instructions.

### Inductively coupled plasma mass spectrometry (ICP-MS)

ICP-MS was used to measure platinum content in biological samples. Cisplatin contains a platinum atom at its core, and therefore measurement of platinum is an indication of cisplatin content. The cochlear tissues were microdissected into stria vascularis, spiral ligament, organ of Corti, and SGNs; vestibular organs were microdissected into utricle, saccule, and cristae. Dissected tissues were submerged in UltraPure™ Distilled Water (10977023, Invitrogen, Waltham, MA) during dissection. Liquid was removed via a speed vacuum concentrator (Eppendorf Vacufuge) and kept frozen at −80°C for Experiment 1 samples and kept at room temperature for Experiment 2 samples until ready for analysis.

As previously mentioned, during mouse tissue harvest, left kidneys were cut in the horizontal plane into three pieces, frozen immediately in dry ice, and stored at −80°C. The middle piece of the fresh frozen kidneys, which included the renal artery and vein, was immersed in 4% PFA (diluted with 1X DPBS from 16% PFA), and fixed overnight at 4°C. Subsequently, kidney samples were washed three times in 1X DPBS, dabbed on a paper towel to absorb excess fluid, and transferred to new centrifuge tubes for sample submission.

ICP-MS was performed at the Mass Spectrometry Core Facility at the University of Massachusetts Amherst using NexION 350D ICP-MS (Perkin-Elmer, Waltham, MA). Organs were digested in 50μl (cochlear tissues) or 100μl (kidney tissues) trace metal grade nitric acid (HNO_3_) and incubated for 20 minutes at 65°C. An equal volume of hydrogen peroxide (Optima Grade, Fisher Chemical) was added, and the incubation was repeated for 20 minutes at 65°C. Samples were spun down for 1 minutes at 14,000xg and diluted 1:20 with MilliQ H_2_O prior to ICP-MS measurements. Platinum (Pt; MW. 195) and sulfur monoxide (SO; MW. 48) were measured in dynamic reaction mode (DRC) using oxygen at a 1.2ml/minutes gas flow rate. Platinum (0.5ppt to 10ppb standards) and sulfur (50 ppb to 500ppb standards) standard curves were generated using single element standards (PerkinElmer) to quantitatively measure levels of platinum and sulfur. Platinum levels were normalized to sulfur levels for each sample.

### Flow cytometry

Spleens were collected, cut in smaller pieces, and placed in media containing RPMI (112-025-101, Quality Biologicals), 5% heat-inactivated FBS (10082-147, Thermo Fisher), and 0.5% Penicillin (P3032, Sigma). Tissues were mechanically disrupted with a 500ul syringe plunger and passed through a 40μm cell strainers (BD Falcon, San Jose, CA, USA) to obtain single cell suspension. Red blood cell (RBC) lysis was performed at room temperature for 5 minutes in 1X RBC lysis buffer (10X RBC lysis buffer diluted in mqH_2_O, 420301, BioLegend) and reaction was stopped by diluting the 1X RBC lysis buffer with 35ml 1X DPBS. Cells were resuspended in FACS buffer (1X DPBS, 5% heat-inactivated FBS, 2mM EDTA) and counted (Cellometer K2, Nexcelom, Lawrence MA). Spleen single cell suspensions were pre-treated with anti-mouse CD16/CD32 antibody (101319, BioLegend) for blocking of Fc receptors followed by surface staining with antibodies in FACS buffer for 30 minutes at 4 °C. Information on the antibodies is presented in Table 1. Cells were then washed and incubated in Fixable Viability dye eFluor™450 (65-0863, eBioscience™) at a 1:1000 dilution in 1X DPBS and incubated for 30 minutes at 4°C. Cells were washed again and flow cytometric acquisition was performed on a BD LSRFortessa (BD Biosciences, Franklin Lakes, NJ). Data were analyzed using FlowJo Software version 10 (BD Biosciences, Ashland, OR). Cells were initially gated on forward scatter (FCS), side scatter (SSC), and viability dye to eliminate cell debris/dead cells and gate on live cell populations. Live cells were gated for CD45 expression, a pan-leukocyte marker. Cell surface markers were used to identify specific populations of immune cells, including macrophages (CD11b+, CD11b+ CX3CR1^GFP^+, and CD11b+ CX3CR1^GFP^-), neutrophils (CD11b+ F4/80-Ly6G+), NK cells (CD3e-NK1.1+), T cells (CD11b-CD3e+), and B cells (CD11b-CD19+).

### Statistics

Statistical analyses were carried out using Graph Pad Prism 9 (GraphPad, San Diego, CA). The Shiapiro-Wilks normality test was employed to verify the normal distribution of the data. A Student’s t-test was performed to compare two groups. Comparisons of multiple groups were conducted using either ordinary one-way ANOVA followed by Tukey’s post-hoc multiple comparisons test or two-way ANOVA to detect the main effect of PLX3397 treatment across multiple groups.

For statistical analysis to assess the protective effect of PLX3397 on male and female mice in Experiment 1, Mixed Effect Modeling was performed in R (version 4.1.3) with the “lmerTest” package. Confidence intervals and p-values were calculated using the t-distribution with degrees of freedom determined via Satterthwaite’s method. Incorporating “animal” as a random effect in the model assumes that each mouse responds differently to the treatment. To evaluate differences in the protection provided by PLX3397 treatment between males and females, an interaction term “Treatment * Sex” was introduced, and the following model was used:

DPOAE = Level + Treatment + Sex + Treatment * Sex + (1 | Animal).

Data are presented as mean ± standard deviation (SD) or mean ± standard error of the mean (SEM). Statistical significance is indicated as: (∗) P < 0.05, (∗∗) P < 0.01, (∗∗∗) P < 0.001, (∗∗∗∗) P < 0.0001. P-value above 0.05 (P>0.05) was considered non-significant and is denoted as “ns”. Statistical comparisons are presented in color-coded remarks (n.s. or asterisks). Saline+Vehicle group is compared to Saline+PLX3397 (purple remarks), Cisplatin+Vehicle (red remarks), and Cisplatin+PLX3397 (green remarks). Comparisons between the Cisplatin+Vehicle and Cisplatin+PLX3397 groups are denoted in black remarks.

## Results

### Development of the cochlear macrophage ablation protocol

Macrophages are the primary resident immune cells in the cochlea and play important roles in the response to cochlear injury (*27, 34, 84–87*). To investigate the roles of macrophages in cisplatin-induced ototoxicity, macrophages were ablated by pharmacologically inhibiting CSF1R signaling. Tissue-resident macrophages and microglia are highly dependent on CSF1R signaling for their survival (*61–63*). PLX3397 is a CSF1R antagonist and an inhibitor for c-Kit and FLT3 that penetrates the BBB and results in elimination of 95-99% of all microglia in the CNS (*61, 63–65, 88*). The discontinuation of PLX3397 leads to robust repopulation of brain microglia within a seven-day period (*62, 63*). While PLX3397 administration via chow feeding effectively ablates cochlear macrophages, mice receiving cisplatin demonstrate decreased appetite, potentially resulting in insufficient drug administration. To account for this, mice were fed with control or PLX3397-formulated chow for seven days prior to cisplatin treatment followed by 50mg/kg PLX3397 administration via oral gavage during the cisplatin administration protocol and until mice were euthanized. Initially, PLX3397 was administered by oral gavage once every three days (Figure 1A). This experimental design will be referred to as “Experiment 1”. The inner ears were examined at 9 days, 14 days, and 28 days after the first cisplatin injection. At day 9, macrophages were depleted; however, by day 14, some macrophage repopulation was observed (Supplemental Figure 1). Therefore, we adjusted the PLX3397 administration frequency, starting on day 22 of the 42-day protocol, from once every three days to daily treatment throughout the remainder of the cisplatin administration protocol. At day 28, following five consecutive days of daily PLX3397 administration, a significant and sustained reduction in cochlear macrophage population was observed (Supplemental Figure 1). At the end of the experiment, mice that received PLX3397 exhibited depletion of approximately 93% of all CX3CR1^GFP^-labeled resident macrophages compared to saline/vehicle-treated mice (Figure 1C-D, Supplemental Figure 1). It is also important to note that administration of cisplatin alone (cisplatin/vehicle) resulted in a 42.15% reduction in the number of cochlear macrophages compared to saline/vehicle-treated mice (Figure 1C-D). Saline-treated mice continued to gain weight throughout the experimental protocol, while all cisplatin-treated mice lost weight. Notably, mice treated with cisplatin/PLX3397 showed significantly reduced weight loss when compared to those treated with cisplatin/vehicle (Figure 1B).

To examine whether macrophages contributed to cisplatin-induced hearing loss, ABRs were measured in control mice and in mice subjected to macrophage ablation via PLX3397. The auditory tests were performed prior to PLX3397 pretreatment (baseline) and at the end of the cisplatin administration protocol (endpoint). Differences in ABR thresholds between the endpoint and baseline measurements were calculated and reported as ABR threshold shifts. In the absence of cisplatin, PLX3397 treatment resulted in no significant changes in hearing sensitivity (Figure 2A-C). Consistent with our previous results (*14, 60*), all cisplatin-treated mice exhibited significant increases in ABR thresholds compared to baseline. Notably, mice treated with both cisplatin and PLX3397 exhibited significantly reduced ABR threshold shifts compared to mice treated with cisplatin alone (cisplatin/vehicle) (Figure 2A). The protective effect of PLX3397 was observed in both cisplatin-treated female and male mice, with greater protection observed in male mice (Figure 2B-C).

**Figure 2.**
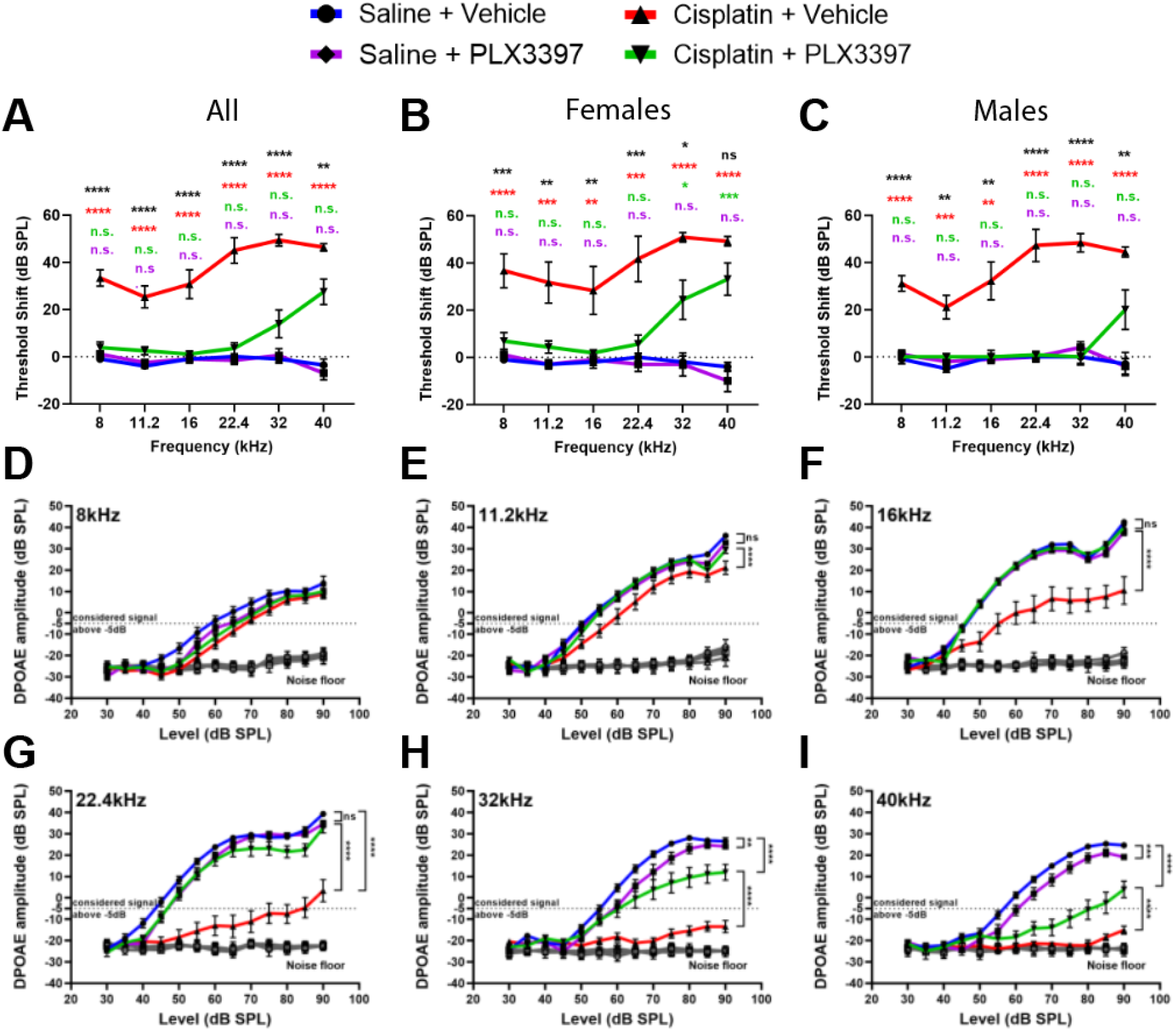
Partial macrophage ablation protected against cisplatin-induced hearing loss and OHC dysfunction (Experiment 1). (A-C) Auditory sensitivity was measured via ABRs at baseline and endpoint. Threshold shifts are reported as the difference between baseline and endpoint ABR thresholds. (A) Cisplatin/vehicle-treated mice (red line) demonstrated significant hearing loss (threshold shifts) at all frequencies compared to saline/vehicle-treated mice (blue line). PLX3397 alone (purple line) did not cause hearing loss. PLX3397 provided significant protection against cisplatin-induced hearing loss (green line). (B-C) PLX3397 protected against cisplatin-induced hearing loss in both (B) female and (C) male mice, with better protection in males than in female (*p=0.0277). *P values were calculated using one-way ANOVA with Tukey’s multiple comparisons test. Mean±SEM, n=10-16 mice per experimental group. Statistical comparisons (asterisks or n.s.) are color-coded as described in Methods*. (D-I) OHC function was measured via DPOAEs. An emission at 2f_1_-f_2_ was considered present when its amplitude was greater than −5dB (dotted line). Mice co-treated with saline and PLX3397 maintained normal DPOAE amplitudes across an f_2_ range of 8-22.4kHz, with a small but significant decrease at 32 and 40kHz compared to control mice (saline/vehicle-treated mice). Mice treated with cisplatin/vehicle showed a significant reduction in DPOAE amplitudes. Macrophage ablation via PLX3397 treatment significantly protected against cisplatin-induced loss of DPOAE amplitudes. *Statistical analyses were performed using two way-ANOVA with Tukey’s multiple comparisons test (main column effect). Mean±SEM, n=10-16 mice per experimental group*.

Since cisplatin treatment leads to death of cochlear outer hair cells (OHCs) (*60, 89*), we next measured DPOAEs, indirectly measuring OHC function (*60, 89, 90*). Mice treated with cisplatin alone (cisplatin/vehicle) demonstrated significant reductions in DPOAE amplitudes at all frequencies (8-40kHz) (Figure 2D-I). Interestingly, mice treated with cisplatin/PLX3397 showed higher DPOAE amplitudes compared to mice treated with cisplatin alone (cisplatin/vehicle), suggesting that macrophage ablation protected against cisplatin-induced OHC dysfunction (Figure 2D-I). While macrophage ablation by PLX3397 protected against cisplatin-induced OHC dysfunction in both male and female mice, we observed a sex difference in the protective effect of PLX3397 against cisplatin-induced OHC dysfunction with better protection in male mice (Supplemental Figure 2).

After the endpoint auditory tests, mice were euthanized, and inner ear tissues were harvested. Cochlear wholemounts were stained for Myosin7a to visualize inner hair cells (IHCs) and OHCs. Cochleae from control mice (saline/vehicle-treated mice) showed normal cochlear morphology with three rows of OHCs and a single row of IHCs (Figure 3A). Mice treated with PLX3397 alone did not show any hair cell loss. Cochleae of cisplatin-treated mice showed significant loss of OHCs at cochlear regions corresponding to frequencies of 22.4 kHz and higher. In contrast, mice co-treated with cisplatin and PLX3397 showed greater OHC survival in these high-frequency regions compared to those treated with cisplatin alone (cisplatin/vehicle) (Figure 3B).

**Figure 3.**
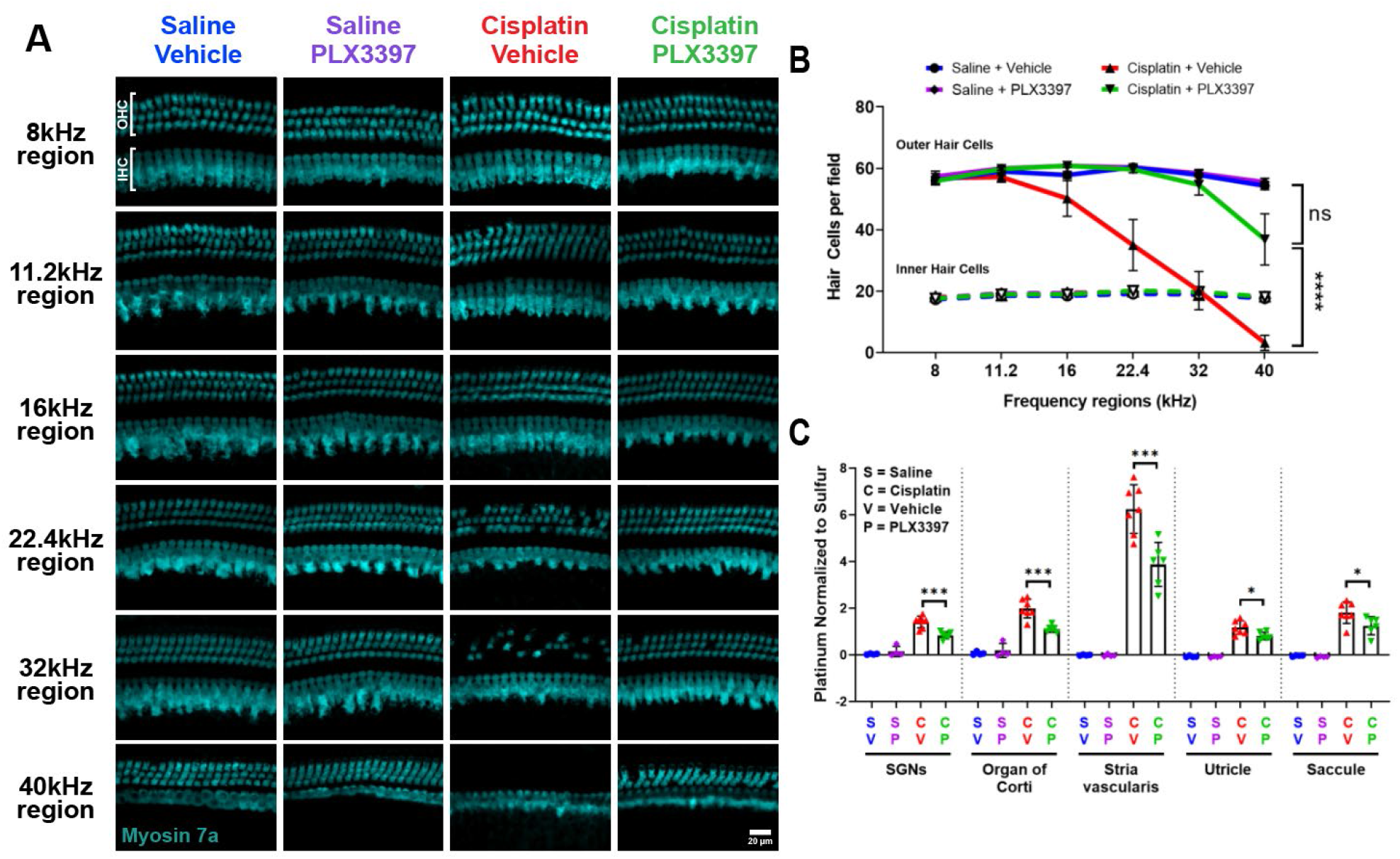
Partial macrophage depletion via PLX3397 increased OHC survival and decreased cisplatin accumulation in the cochleae of cisplatin-treated mice (Experiment 1). (A-B) PLX3397 alone did not affect the number of OHCs. Cisplatin/vehicle-treated mice show a significant reduction in the number of OHCs in the higher frequency regions (22.4 kHz and above). PLX3397 protected against cisplatin-induced OHC death. The number of IHCs was not different in any of the four treatment groups. (A) *Scale bar, 20 μm.* (B) Quantification of OHC and IHC numbers. The solid lines (blue, purple, red, green) represent OHC counts, and dotted lines represent IHC counts. *Mean±SEM, n=6 cochleae per experimental group. Statistical analysis was performed using two way-ANOVA with Tukey’s multiple comparisons test (main column effect)*. (C) Platinum concentrations in microdissected inner ear tissues were measured using ICP-MS. Cochleae of control (no cisplatin) mice do not contain measurable platinum. Cochleae from mice treated with cisplatin showed significant increases in platinum in all cochlear regions. Mice treated with cisplatin and PLX3397 exhibited significantly lower platinum concentrations in all cochlear tissues, with stria vascularis showing the largest difference. *Mean±SD, n=4-7 inner ears per experimental group. Statistical analysis was performed using one way-ANOVA with Tukey’s multiple comparisons test*.

Cisplatin has been shown to cause damage to SGNs (*91*) in addition to hair cells (*60, 89*). To investigate the effects of cisplatin and macrophage ablation by PLX3397 on SGNs, cochlear sections were immunostained for the neuronal marker Tuj-1. Quantitative analyses of three regions of the cochlea (apex, middle, and base) showed that cisplatin resulted in decreased SGN density at the apex, but not in the middle or basal cochlear regions. Co-treatment with cisplatin and PLX3397 resulted in preservation of SGN cell bodies in the apical region of the cochlea (Supplemental Figure 3). Together, these data indicate that macrophage ablation was protective against cisplatin-induced hearing loss, OHC death, and SGN loss.

Previously we demonstrated that cisplatin enters the cochlea within one hour after systemic administration and is retained indefinitely in both mouse and human cochleae (*14*). To elucidate the underlying mechanisms by which macrophage ablation by PLX3397 confers protection against cisplatin-induced ototoxicity, we examined whether PLX3397 alters cochlear uptake or retention of cisplatin. Platinum levels in microdissected cochlear (SGNs, organ of Corti, and stria vascularis) and vestibular tissues (utricle and saccule) of the inner ear were measured using ICP-MS. Consistent with our previous results (*14*), platinum levels were significantly increased in the inner ears of cisplatin-treated mice, with the highest platinum accumulation observed in the stria vascularis (Figure 3C). Mice co-treated with cisplatin and PLX3397 showed significantly less platinum accumulation in all microdissected inner ear tissues. Notably, the most substantial difference was observed in the stria vascularis, which showed a 33% reduction in cisplatin accumulation in PLX3397-treated animals compared to those treated with cisplatin alone (cisplatin/vehicle) (Figure 3C). These data indicate that the protective effect of macrophage ablation by PLX3397 is likely related to a reduction in cisplatin accumulation in the inner ear.

### Complete and sustained macrophage ablation fully protects against cisplatin-induced ototoxicity

In the study presented above and in Figures 1-3 (Experiment 1), we showed that macrophage ablation using PLX3397 reduces cisplatin-induced hearing loss and inner ear damage. However, the PLX3397 treatment regimen used in Experiment 1 was insufficient to achieve complete macrophage ablation, as demonstrated by the partial macrophage repopulation observed (Supplemental figure 1). To address this limitation and to test the effects of more complete macrophage ablation, we repeated the study with a modified experimental design wherein mice were fed with control or PLX3397-formulated chow for seven days prior to cisplatin treatment, followed by daily 50mg/kg PLX3397 administration via oral gavage throughout the cisplatin administration protocol (Experiment 2, Figure 4A). Mice that received PLX3397 via chow for seven days followed by daily oral gavage effectively ablated 96% of all CX3CR1^GFP^-positive macrophages within cochlear sections, compared to saline/vehicle-treated mice (Figure 4C-D). Furthermore, we extended our analysis to visualize CX3CR1^GFP^-positive macrophages from cochlear and stria vascularis wholemounts to comprehensively evaluate the extent of macrophage ablation by PLX3397. Quantification of CX3CR1^GFP^-positive macrophages in both tissues revealed an ablation efficiency of >88% in mice administered PLX3397 (Supplemental Figure 4-1), without affecting the peripheral immune cells in the spleen, including macrophages, T cells, B cells, NK cells, and neutrophils (Supplemental figure 4-2). Similar to Experiment 1, the treatment with cisplatin alone (cisplatin/vehicle) again resulted in a 20-25% reduction in cochlear macrophages, specifically in the modiolus, osseous spiral lamina, and the stria vascularis (Figure 4C-D, Supplemental Figure 4-1).

**Figure 4.**
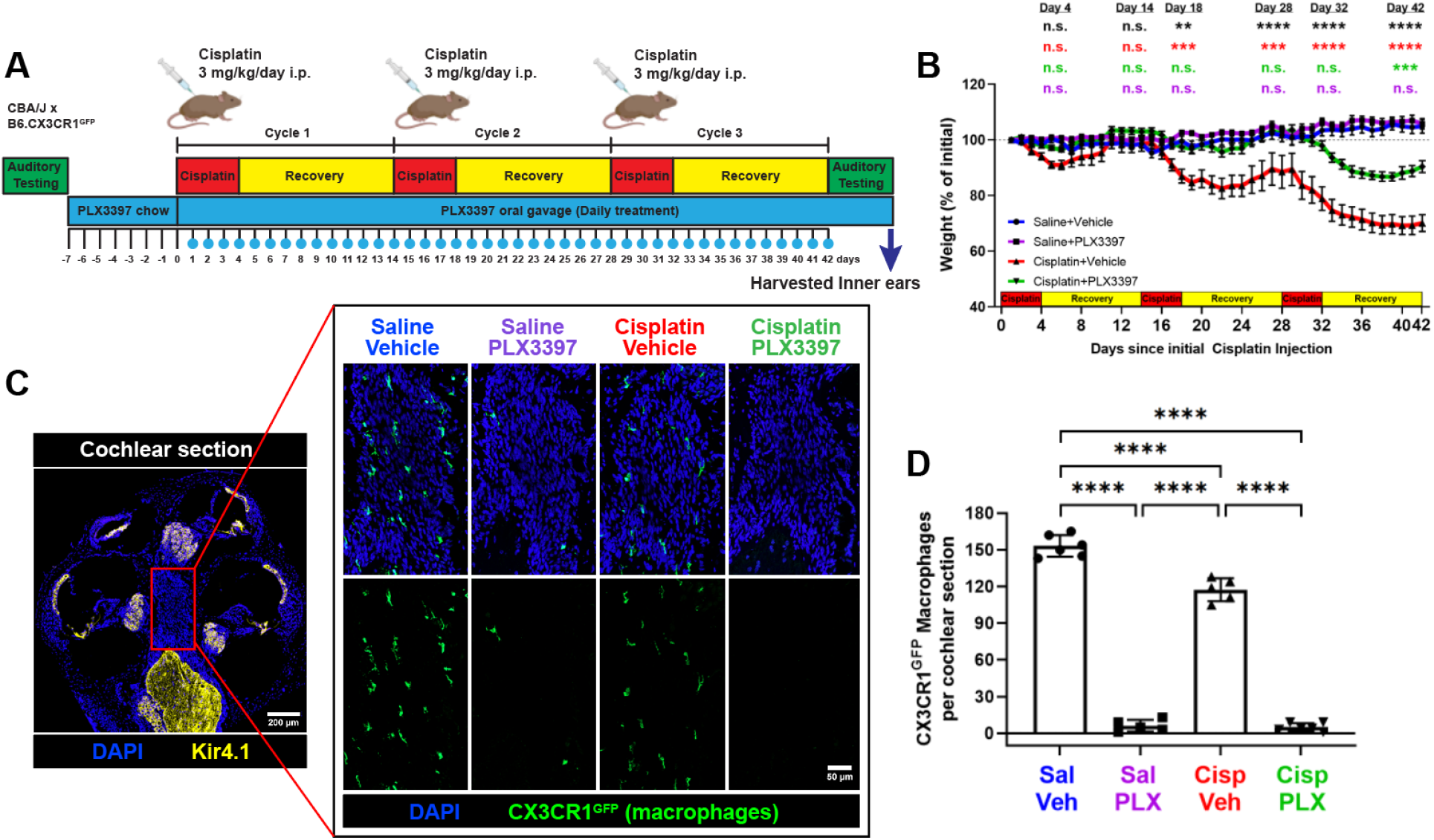
PLX3397 in rodent chow followed by daily oral gavage resulted in sustained macrophage ablation and a significant reduction in weight loss induced by cisplatin (Experiment 2). (A) Experimental design of Experiment 2. Mice received either control chow or PLX3397-formulated chow for seven days to facilitate macrophage depletion. Following initiation of cisplatin administration, mice received daily PLX3397 treatment via oral gavage, which continued until euthanasia, ensuring sustained macrophage ablation. The days on which mice received PLX3397 through oral gavage are indicated with blue circles. (B) Mice co-administered cisplatin and vehicle exhibited significant weight loss, whereas those co-administered cisplatin and PLX3397 demonstrated significantly reduced cisplatin-induced weight loss. *Mean±SEM, n=8-9 mice per experimental group. Statistical analysis was performed using one-way ANOVA with Tukey’s multiple comparisons test. Statistical comparisons (asterisks or n.s.) are color-coded as described in Methods*. (C-D) Following sustained macrophage ablation via PLX3397, more than 96% of all macrophages were ablated in cochleae collected after endpoint auditory testing when compared to saline/vehicle-treated mice. (C) *Scale bars in 200 μm (cochlear section) and 50 μm (modiolus, magnified inset).* (D) Quantification of macrophages in whole cochlear sections. *Mean±SD, n=5-6 cochleae per experimental group. Statistical analysis was performed using one-way ANOVA with Tukey’s multiple comparisons test*.

Together these data demonstrate that CSF1R inhibition via PLX3397 chow followed by daily oral gavage leads to sustained depletion of resident cochlear macrophages while minimally affecting the peripheral immune cells in both saline- and cisplatin-treated mice.

Throughout Experiment 2, mice were weighed daily and weights are reported as a percentage of initial body weight on day 1. Mice in both saline/vehicle- and saline/PLX3397-treated groups exhibited modest weight gain throughout the study. While all cisplatin-treated mice demonstrated significant weight loss, mice co-treated with cisplatin and PLX3397 exhibited significantly less weight loss compared to those treated with cisplatin alone (cisplatin/vehicle) (Figure 4B). To determine whether sustained macrophage ablation by PLX3397 protected against cisplatin-induced hearing loss in Experiment 2, we again measured ABRs and DPOAEs. Mice treated with cisplatin alone (cisplatin/vehicle) had significant hearing loss (ABR threshold shifts) and substantial reduction in DPOAE amplitudes at all frequencies tested (Figure 5). Remarkably, mice co-administered cisplatin and PLX3397 displayed virtually no ABR threshold shifts and complete protection of OHC function, suggesting that PLX3397 treatment provided complete protection against cisplatin-induced hearing loss (Figure 5A). This complete protection was observed in both male and female animals (Figure 5B-C; Supplemental Figure 5). Moreover, mice treated with PLX3397 also demonstrated near-total protection against cisplatin-induced OHC death (Figure 6A-B). Taken together, these data indicate that sustained macrophage ablation by PLX3397 provides complete protection against cisplatin-induced hearing loss, OHC dysfunction, and OHC death.

**Figure 5.**
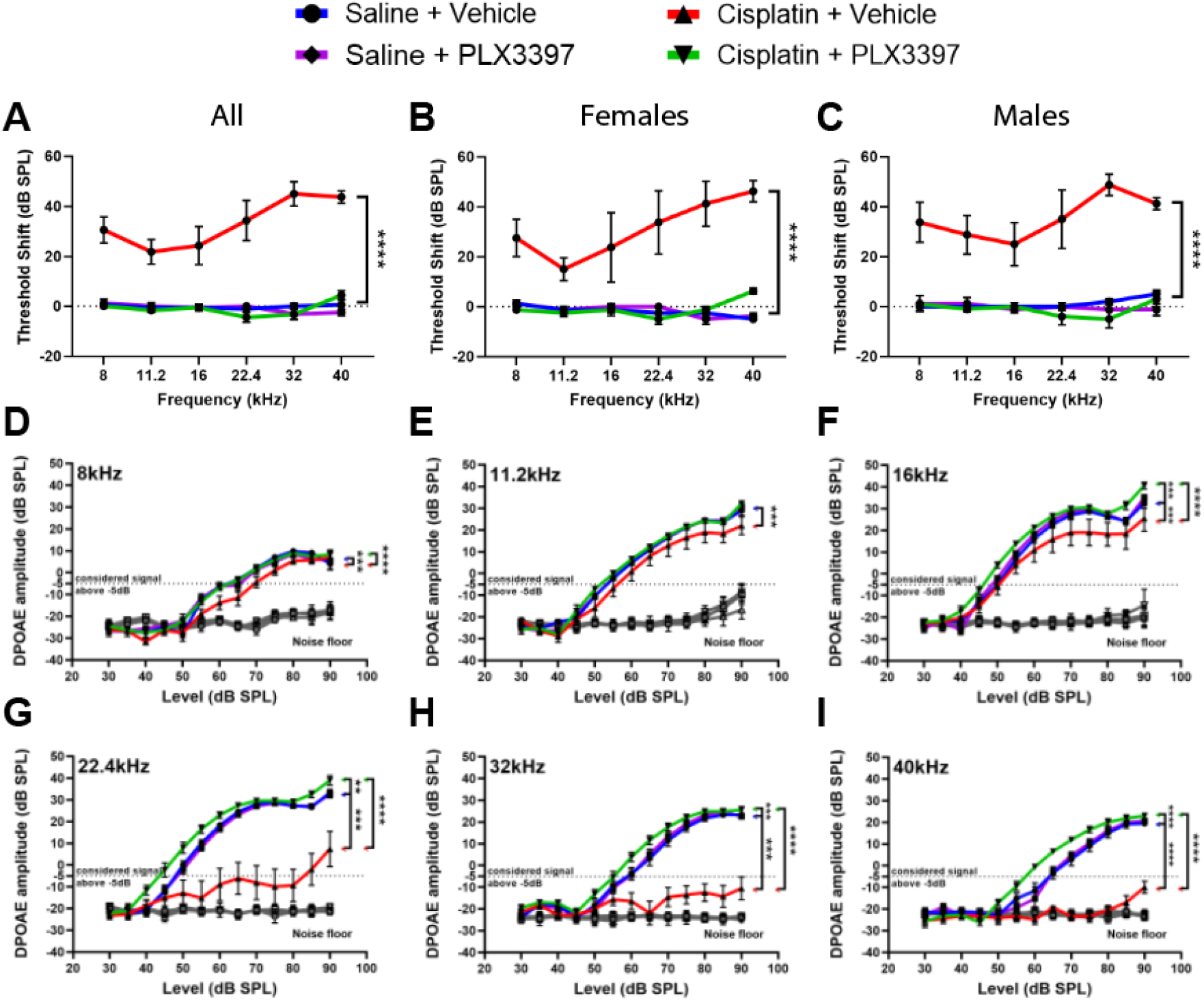
Sustained depletion of macrophages via PLX3397 provided complete protection against cisplatin-induced hearing loss and OHC dysfunction (Experiment 2). Hearing loss was assessed by ABRs, and OHC function was evaluated using DPOAEs. (A-C) Auditory tests were performed before (baseline) PLX3397 treatment and after (endpoint) completion of the cisplatin administration protocol. Hearing loss is reported as threshold shifts (the difference between baseline and endpoint ABR thresholds). PLX3397 in saline-treated mice (purple line) did not cause hearing loss when compared to saline/vehicle-treated mice (blue line). Cisplatin/vehicle-treated mice (red line) exhibited significant hearing loss (threshold shifts) at all frequencies. Cisplatin-treated mice receiving PLX3397 showed comparable ABR threshold shifts to saline-treated mice, indicating that PLX3397 resulted in complete protection against cisplatin-induced hearing loss. This protection was observed in both (B) female and (C) male mice. (D-I) PLX3397 in saline-treated mice did not elicit significant changes in OHC function (DPOAE amplitudes). Cisplatin/vehicle-treated mice (red line) exhibited significantly reduced DPOAE amplitudes compared to saline-treated mice. PLX3397 resulted in complete protection against cisplatin-induced OHC dysfunction. *An emission at 2f_1_-f_2_ was considered present when its amplitude exceeded the threshold of −5dB (dotted line). The grey line represents the biological noise floor. For both ABRs and DPOAEs, data are shown as Mean±SEM, n=8-9 mice per experimental group. Statistical analysis was performed using two way-ANOVA with Tukey’s multiple comparisons test (main column effect)*.

**Figure 6.**
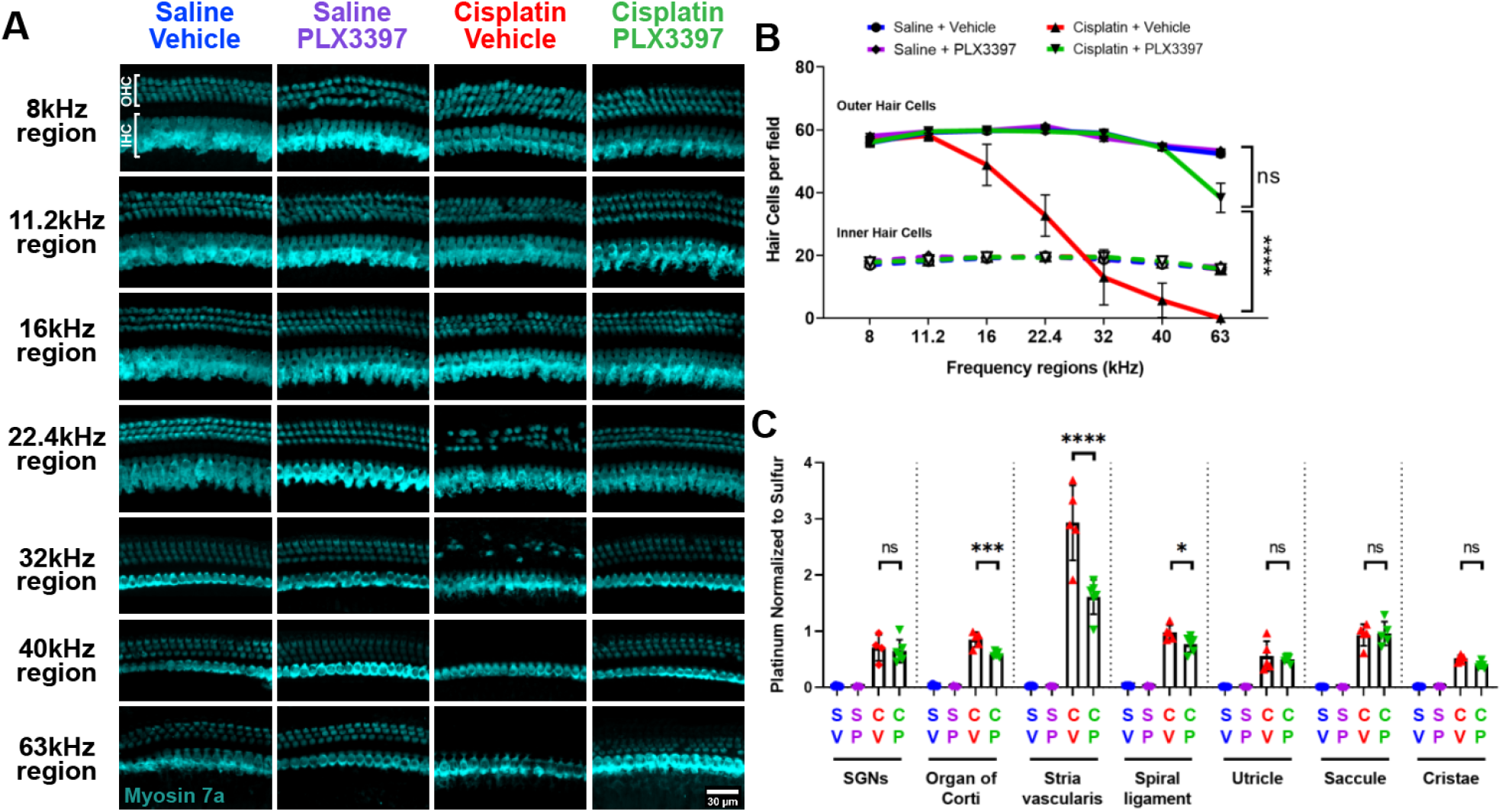
Sustained macrophage ablation protected against cisplatin-induced OHC death and resulted in reduced cisplatin accumulation in the cochlea (Experiment 2). (A-B) Cochlear wholemounts were stained for myosin 7a (cyan) to visualize hair cells. (A) Representative images and (B) quantitative analysis revealed that cisplatin administration resulted in death of OHCs at the 22.4 kHz region and above. PLX3397 provided near-complete protection against cisplatin-induced OHC loss, resulting in OHC numbers that were comparable to those in saline-treated mice. IHC counts did not differ in any of the experimental groups. *(A) Scale bar, 100 μm. (B) Data are shown as Mean±SEM, n=6 cochleae per experimental group. Statistical analysis was performed using two way-ANOVA with Tukey’s multiple comparisons test (main column effect).* (C) Platinum levels were measured using ICP-MS in microdissected inner ear tissues. Platinum was not detected in cochleae of mice that received saline/vehicle or saline/PLX3397 treatments. A significant increase in platinum was observed in all tissue samples from mice treated with cisplatin/vehicle, with the highest levels observed in the stria vascularis. Mice co-administered with cisplatin and PLX3397 exhibited significantly reduced platinum levels in the organ of Corti and spiral ligament, with the stria vascularis showing the largest reduction. No significant differences in platinum levels were observed in the SGNs or vestibular tissues when compared with mice that received cisplatin/vehicle treatment. *Data are shown as Mean±SD, n=4-6 cochleae per experimental group. Statistical analysis was performed using one-way ANOVA with Tukey’s multiple comparisons test*.

Although cisplatin does not result in the death of IHCs in the cochlea, it does result in loss of specialized ribbon synapses that mediate rapid and sustained release of neurotransmitters from IHCs (*92, 93*). To assess the effects of cisplatin and macrophage ablation on IHC synapses, cochlear wholemounts were stained for both Carboxy-terminal Binding Protein 2 (CtBP2), which labels presynaptic ribbons, and for GluR2, an AMPA receptor subunit that labels postsynaptic glutamate receptors. Numbers of juxtaposed CtBP2-GluR2 puncta (“synapses”) per IHC and CtBP2-positive puncta (“orphan ribbons”) were quantified. PLX3397 treatment alone did not result in any loss of synapses compared to controls (saline/vehicle-treated mice) (Supplemental Figure 6A-B). Mice treated with cisplatin alone (cisplatin/vehicle) demonstrated synaptic loss in the lower frequency (8, 11.2, 16kHz) regions (Supplemental Figure 6A-B). Some animals treated with cisplatin/vehicle displayed an increase in the number of orphan ribbons, characterized by the absence of juxtaposed GluR2 in relation to the CtBP2 (Supplemental Figure 6A,C) (*94*). Ablation of macrophages using PLX3397 in cisplatin-treated mice resulted in complete protection against loss of synapses induced by cisplatin (Supplemental Figure 6A-C). Concurrent with the loss of synapses, quantification of SGN cell bodies revealed a significant decrease in SGN density in the apical cochlear regions in cisplatin-treated mice, while mice that received both cisplatin and PLX3397 exhibited significant protection against cisplatin-induced loss of SGN cell bodies (Supplemental Figure 6D-E). These data demonstrate that sustained macrophage ablation using PLX3397 provided complete protection against cisplatin-induced loss of SGNs and concomitant loss of synapses in the inner ear.

We again examined whether administration of PLX3397 reduced cisplatin accumulation in the inner ear in Experiment 2 using ICP-MS. Platinum was significantly increased in cochlear and vestibular tissues from cisplatin-treated mice relative to saline-treated mice, with the highest accumulation again observed in the stria vascularis. PLX3397 treatment resulted in reduced platinum levels in the stria vascularis, organ of Corti, and spiral ligament (Figure 6C). Overall, sustained macrophage ablation in Experiment 2 led to an average reduction in platinum accumulation of 45.05% in the stria vascularis, compared to the 33.62% reduction observed in Experiment 1 where we observed partial macrophage repopulation. Together, these data suggest a greater reduction in platinum accumulation within the stria vascularis with sustained depletion of macrophages in the cochlea, supporting the idea that the protective effect of macrophage ablation via PLX3397 occurs through a mechanism involving the reduction of cisplatin uptake into the inner ear.

### Macrophage ablation using PLX3397 confers protection against cisplatin-induced nephrotoxicity

Cisplatin causes both ototoxicity and nephrotoxicity. Therefore, we evaluated the effects of macrophage ablation using PLX3397 treatment on cisplatin-induced kidney dysfunction and injury (*60*). To examine this, blood and kidney tissues from Experiment 2 were analyzed. Following the completion of cisplatin administration and auditory tests, plasma blood urea nitrogen (BUN) and neutrophil gelatinase-associated lipocalin (NGAL) levels, biomarkers of kidney function and injury, were measured. Mice treated with cisplatin exhibited significantly elevated levels of plasma BUN and NGAL. However, PLX3397 prevented cisplatin-induced elevation of both BUN and NGAL levels, suggesting that macrophage ablation protected against cisplatin-induced renal dysfunction (Figure 7A-B). Furthermore, histological assessments using Periodic Acid-Schiff (PAS) and Masson’s trichrome (MT) staining revealed a substantial induction of kidney injury and maladaptive repair, characterized by significant tubular injury and interstitial fibrosis, in cisplatin-treated mice. Macrophage ablation using PLX3397 provided significant protection against both cisplatin-induced tubular injury and interstitial fibrosis in the kidney (Figure 7C-F). These data indicate that PLX3397 effectively protected against cisplatin-induced renal tissue damage and associated pathological changes.

**Figure 7.**
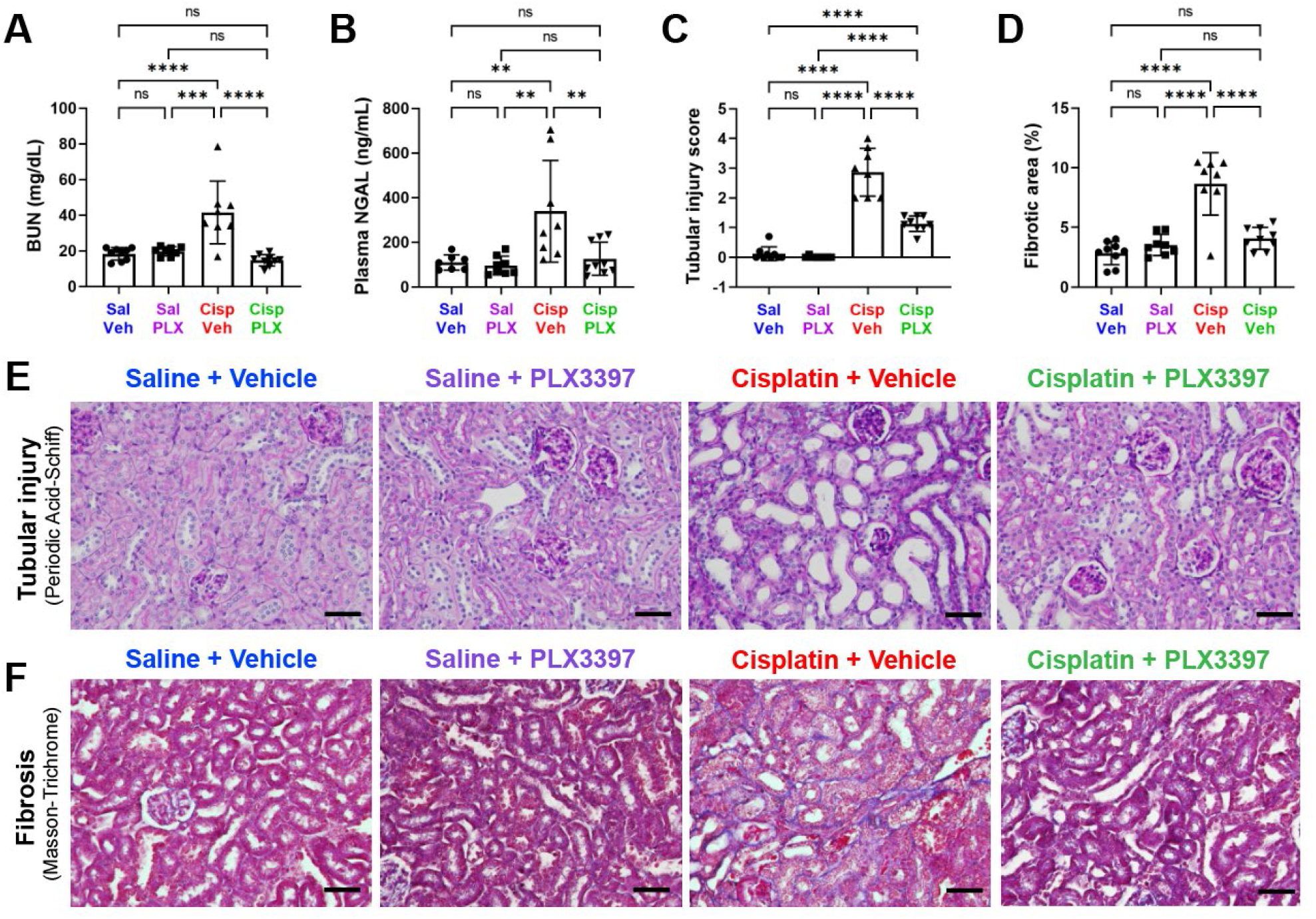
Sustained depletion of macrophages protects against cisplatin-induced kidney damage (Experiment 2). Kidney function, tubular injury, and fibrosis were evaluated after endpoint auditory testing. Macrophage ablation using PLX3397 protected against cisplatin-induced increases in (A) plasma blood urea nitrogen (BUN) levels and (B) Neutrophil gelatinase-associated lipocalin (NGAL) levels. (C) Tubular injury was assessed semi quantitatively using a tubular injury score (see Methods). Cisplatin resulted in increased tubular injury scores, while PLX3397 protected against cisplatin-induced injury. (D) The percentage of fibrotic area in the entire surface area of the section was calculated. PLX3397 protected against cisplatin-induced fibrosis. (E-F) Representative images of cortico-medullary junction are presented, demonstrating (E) tubular injury using periodic acid-Schiff staining and (F) fibrosis using Masson-Trichrome staining. *(A-D) Data are expressed as mean±SD, n=8–9 blood samples kidneys per experimental group. P-values were calculated using one-way ANOVA with Tukey’s multiple comparisons test*. (E-F) *The scale bar indicates 50 µm*.

Administration of PLX3397 has been demonstrated to ablate microglia (*61–63*) and cochlear macrophages (Figure 1, Figure 4, Supplemental Figure 4-1). However, the effects of PLX3397 treatment on kidney resident macrophages had not been evaluated. Therefore, we assessed the impact of PLX3397 on kidney resident macrophages. We observed a significant (2.36-fold) increase in the number of CX3CR1^GFP^-positive macrophages within the kidneys of cisplatin/vehicle-treated mice compared to the kidneys of control mice treated with saline/vehicle (Figure 8A-B). Consistent with our observations in the cochlea, PLX3397 resulted in a marked decrease of 95.6% in saline-treated mice (saline/vehicle vs saline/PLX3397) and a reduction of 92.6% in cisplatin-treated mice (cisplatin/vehicle vs cisplatin/PLX3397) of CX3CR1^GFP^-positive macrophages in the kidney (Figure 8A-B).

**Figure 8.**
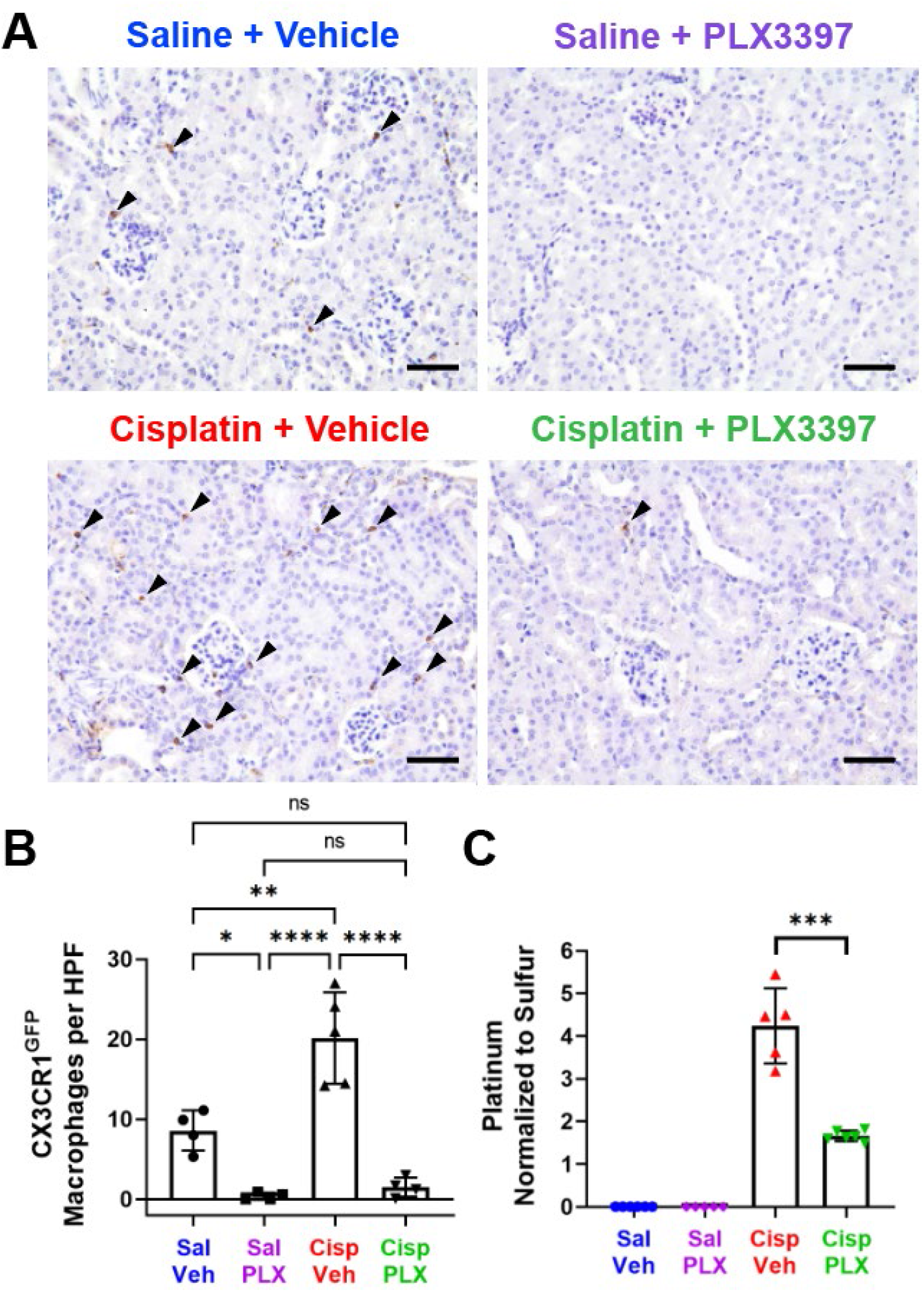
PLX3397 results in ablation of CX3CR1^GFP^-positive cells and significant reduction in cisplatin accumulation in the kidney (Experiment 2). (A) Representative images of renal CX3CR1^GFP^-positive cells (arrowheads), identified through immunohistochemical staining. *Scale bar indicates 50 µm*. (B) The number of CX3CR1^GFP^-positive cells in each field was quantified (n = 4–5 mice per experimental group). Compared to control mice (saline/vehicle-treated mice), PLX3397 treatment in saline-treated mice and cisplatin-treated mice ablated 95.63% and 82.52% of CX3CR1^GFP^-positive cells, respectively. (C) Platinum levels in the kidney tissues were analyzed by ICP-MS (n = 5–6 mice per experimental group). Cisplatin resulted in increased platinum in the kidney, while PLX3397 significantly reduced cisplatin accumulation. *Platinum levels were normalized to sulfur contained in the kidney. Data are expressed as mean±SE. P-values were calculated using one-way ANOVA with Tukey’s multiple comparisons test*.

To begin to examine the mechanisms underlying the protective effects of PLX3397 against cisplatin-induced nephrotoxicity, we investigated whether PLX3397 reduced cisplatin accumulation in the kidney. Platinum levels in kidney tissues were analyzed using ICP-MS. As expected, platinum levels were significantly elevated (4.24-fold) in kidneys from cisplatin/vehicle-treated mice relative to those from saline/vehicle-treated mice. Kidneys that were co-treated with cisplatin and PLX3397 contained 60.88% less platinum compared to kidneys of mice treated with cisplatin/vehicle (Figure 8C). These data indicate that macrophage ablation by PLX3397 protected against cisplatin-induced nephrotoxicity as assessed by biomarkers of kidney function, tubular injury, and interstitial fibrosis, and they suggest that this protective effect is mediated through a mechanism involving reduced cisplatin uptake and/or accumulation in the kidney.

## Discussion

Cochlear macrophages play an important role in homeostasis and during cochlear injury. PVMs are closely associated with the blood vessels and play a pivotal role in regulating the tight junction proteins between endothelial cells that comprise the BLB (*34, 35*). In noise damage models that result in the loss of hair cell synaptic ribbons, macrophages are involved in synaptic repair and the restoration of cochlear synapses and synaptic function (*27*). Here we show that macrophages also play important roles in the cochlear and kidney responses to cisplatin-induced damage, in part by regulating cisplatin uptake and accumulation in the cochlea and kidney.

### Macrophage depletion and cisplatin-induced hearing loss

Our experimental design of macrophage ablation in a mouse model of cisplatin ototoxicity utilized two methods to administer PLX3397. In both Experiment 1 and Experiment 2, the use of PLX3397-formulated chow for 7 days resulted in the ablation of >88% of all cochlear macrophages. However, we observed some macrophage repopulation in Experiment 1, which was ameliorated by increasing the frequency of PLX3397 administration. Data from both studies consistently demonstrated that macrophage ablation conferred significant protection against cisplatin-induced hearing loss, OHC dysfunction and death, loss of SGNs and a concomitant loss of synapses in the inner ear. We have elected to present both studies herein because they collectively demonstrate that the extent of macrophage ablation impacts the magnitude of the protective effect. Notably, Experiment 2, characterized by sustained macrophage ablation, exhibited considerably higher levels of protection approaching near-complete preservation of auditory function and hair cells compared to Experiment 1, in which macrophage repopulation occurred during the cisplatin administration protocol.

The major route of cisplatin entry into the cochlea is thought to be through the BLB in the stria vascularis (*95*). PVMs are closely associated with capillaries and regulate the BLB permeability (*34, 36*) (Figure 9). Treatment with PLX3397 resulted in ablation of PVMs within the stria vascularis (Supplemental Figure 4-1C-D) and reduced platinum levels in the cochlea, with the highest reduction observed in the stria vascularis (Figure 3C; Figure 6C). Thus, our data are consistent with a model in which ablation of PVMs in the stria vascularis resulted in reduced permeability of the BLB and consequently decreased cisplatin entry into the cochlea. The precise cellular and molecular mechanisms by which PVMs regulate BLB permeability during cisplatin treatment remain largely unknown and are likely to involve multiple factors. One proposed mechanism by which the permeability of the BLB may be regulated involves the interaction between PVMs and vascular endothelial cells. PVMs can secrete soluble factors that subsequently bind to their receptors expressed on vascular endothelial cells lining the capillary lumen. This interaction can modulate the expression levels of tight junction proteins in the endothelial cells, thereby influencing BLB permeability (*34, 38, 39*). Therefore, in the context of cisplatin treatment, we propose that soluble factors produced by PVMs may bind to their receptors expressed on vascular endothelial cells, subsequently downregulating tight junction proteins connecting the vascular endothelial cells. This can result in increased permeability of the BLB (leaky BLB), thereby facilitating the entry of cisplatin into the inner ear (Figure 9A). In the absence of PVMs, the soluble factors responsible for BLB breakdown are absent, BLB integrity is maintained, and cisplatin entry into the inner ear is reduced. This model is consistent with a recent report demonstrating that cisplatin-induced hearing loss was associated with increased BLB permeability and reduced expression of tight junction proteins between vascular endothelial cells (*96*). This hypothesis also provides a plausible explanation for the differences in the extent of protection against cisplatin-induced hearing loss that we observed between Experiment 1 and Experiment 2. Our data are consistent with a model in which macrophage repopulation in Experiment 1 may have allowed for the release of soluble factors that disrupted the integrity of the BLB, increasing its permeability and thus increasing the entry of cisplatin into the cochlea. In Experiment 2, sustained ablation of macrophages ensured the maintenance of BLB integrity, thereby limiting the entry of cisplatin into the cochlea. We observed a significant reduction in cochlear platinum levels following macrophage ablation by PLX3397 in both Experiment 1 and Experiment 2, with the largest reduction in the stria vascularis. In Experiment 1, in which partial macrophage repopulation was observed, platinum accumulation in the stria vascularis was reduced by 33.62% in PLX3397-treated animals. In Experiment 2 there was sustained macrophage ablation, and we observed a larger 45.04% decrease in cisplatin accumulation (Figure 3C; Figure 6C). These data suggest that macrophage repopulation in Experiment 1 was associated with greater cisplatin uptake and accumulation in the cochlea compared to Experiment 2, in which macrophage ablation was sustained.

**Figure 9.**
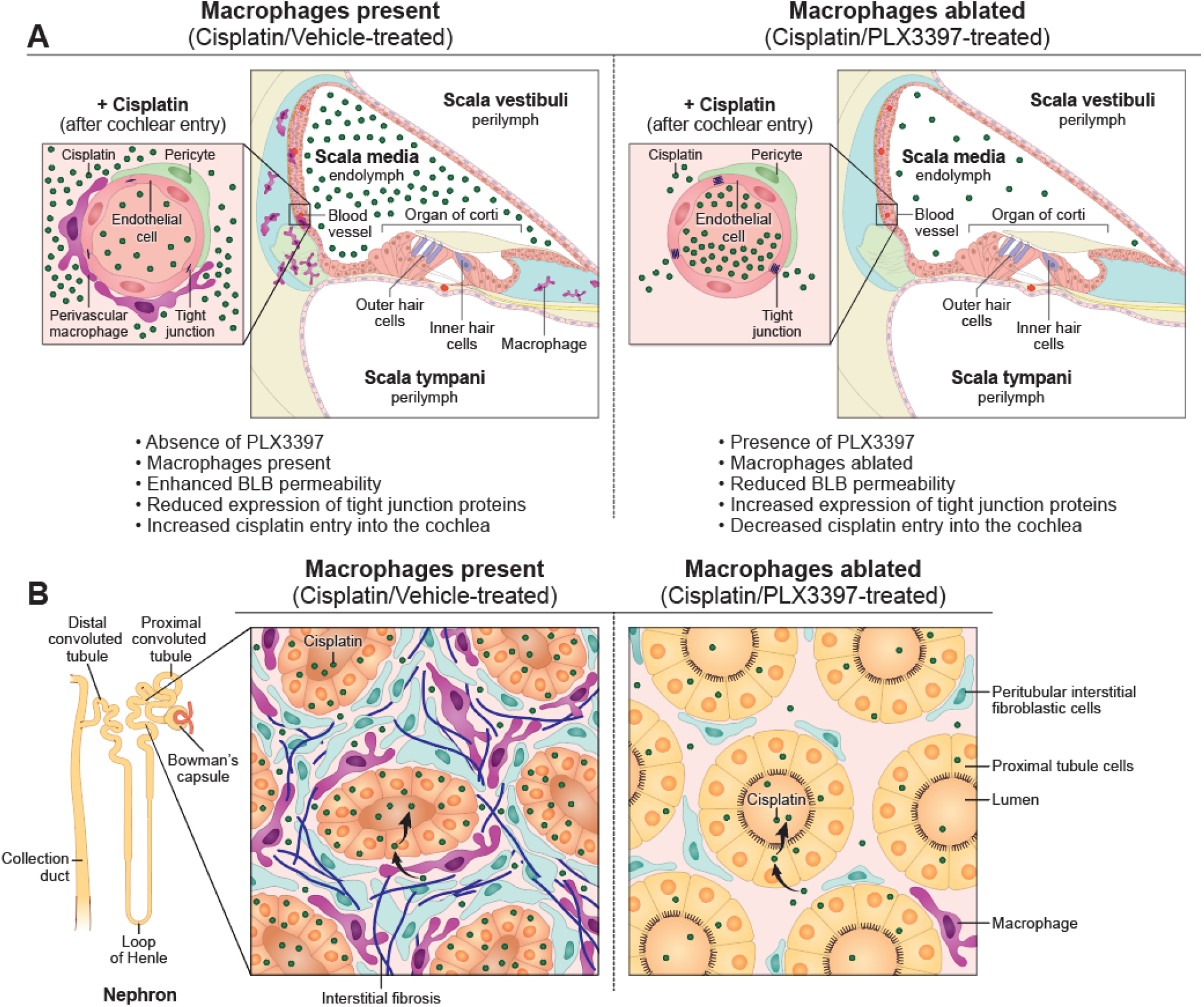
Working model of the mechanisms underlying the protective effects of macrophage ablation against cisplatin-induced hearing loss and kidney injury. (A) In the absence of PLX3397, macrophages are present, and the cochlear BLB is permeable enough to permit cisplatin to cross the BLB and enter the cochlea, where it results in the death of OHCs, loss of ribbon synapses and SGNs, and hearing loss. In the presence of PLX3397, perivascular macrophages are ablated, and cisplatin entry into the cochlea is reduced, resulting in protection against cisplatin-induced hearing loss. (B) In the absence of PLX3397, renal macrophages are present and may promote cisplatin entry into proximal tubule cells causing renal fibrosis and kidney dysfunction. In the presence of PLX3397, macrophages are ablated in the kidney and cisplatin accumulation in the kidney is reduced, thus providing protection against cisplatin-induced tubular injury and interstitial fibrosis.

Although the observed difference in the platinum levels between the two studies suggests a role for macrophages in regulating cisplatin entry into the cochlea, this likely does not fully account for the different levels of observed protection between the two studies. Our analysis of other microdissected inner ear tissues (SGNs, organ of Corti, spiral ligament, utricle, saccule, cristae), revealed minimal differences in platinum levels between the two studies following macrophage ablation in cisplatin-treated mice, in comparison to mice with intact macrophages (Figure 3C; Figure 6C). Thus, additional macrophage-dependent factors may contribute to cisplatin-induced ototoxicity. Cisplatin ototoxicity is associated with increased production of reactive oxygen species generated through NADPH oxidase 3 (*97, 98*) as well as secretion of proinflammatory cytokines such as TNFα, IL-1β, and IL-6 (*99, 100*) through activation of NF-κB, a transcription factor and regulator of innate immunity (*101*). Both oxidative stress and proinflammatory cytokines exert cytotoxic effects in the context of cisplatin-induced ototoxicity (*102–104*). It is plausible that macrophages may also contribute to the production of these cytotoxic factors. Emerging technologies such as single-cell resolution spatial transcriptomics may provide insight into the functional state of macrophages in response to cisplatin exposure in the inner ear.

Upon entering the stria vascularis from the BLB, cisplatin subsequently enters the endolymph within the cochlear duct (*105–108*). Once inside the cochlea, cisplatin can enter various cell types, including hair cells, SGNs, and stria vascularis, via the mammalian cation transporters or possibly via mechanoelectrical transduction channels (*109*) and can exert cytotoxic effects at the level of individual cells (*14, 90, 110–115*). Following systemic administration, cisplatin is present in the organ of Corti, SGNs, stria vascularis, spiral ligament, utricle, saccule, and cristae (Figure 3C, Figure 6C) (*14*). Upon entry into the OHCs, cisplatin irreversibly binds to cellular macromolecules (DNA, RNA, and proteins) and leads to apoptotic degeneration of OHCs and permanent hearing loss (*1*). In contrast to OHCs, IHCs are relatively resistant to the cytotoxic effect of cisplatin, as they appear morphologically intact after cisplatin treatment (Figure 3A-B, Figure 6A-B) (*60, 89*). However, our data demonstrate a significant loss of IHC synaptic ribbons (Supplemental Figure 6A-C) and a reduction in SGN density (Supplemental Figure 3, Supplemental Figure 6D-E) following cisplatin treatment. Interestingly, our data revealed a selective loss of IHC synapses in the low-frequency, apical regions of the cochlea upon cisplatin treatment, while the degeneration of OHCs primarily occurred in the high-frequency, basal region of the cochlea. This loss of IHC synapses in the apical region of the cochlea is congruent with the loss of SGN density in the apical regions, while the basal region remained unaffected (Supplemental Figure 3, Supplemental Figure 6D-E). These observations demonstrate that different cell types within the inner ear exhibit distinct susceptibility profiles in response to cisplatin or other cochlear insults, possibly influenced by tonotopic regions of the organ of Corti or specific organs within the inner ear. For example, basal OHCs are more vulnerable to a variety of cochlear insults, including cisplatin (*60*), gentamicin (*116, 117*), and neomycin (*118*) compared to apical OHCs. Remarkably, macrophage ablation by PLX3397 treatment in both Experiment 1 and Experiment 2 resulted in significant protection against cisplatin-induced loss of IHC synaptic ribbons and SGN cell bodies (Supplemental Figure 3; Supplemental Figure 6). These data provide additional evidence in support of the hypothesis that the presence of PVMs increases BLB permeability, leading to increased cisplatin entry into the cochlea, while macrophage ablation maintains BLB integrity, thereby reducing entry of cisplatin into the cochlea and offering protection against cisplatin-induced cellular damage in the inner ear. Thus, our data suggest a crucial role for PVMs in modulating BLB permeability, highlighting their potential as therapeutic targets for mitigating cisplatin ototoxicity.

### Macrophage depletion and cisplatin-induced nephrotoxicity

Nephrotoxicity is another significant and commonly observed side effect of cisplatin. Therefore, we examined the effects of macrophage ablation on cisplatin-induced nephrotoxicity. We began by examining the nephrotoxic effects of cisplatin in our mouse model, which had been developed for ototoxicity (*60*). Consistent with observations in other mouse models of cisplatin-induced nephrotoxicity (*83, 119–124*), our three-cycle cisplatin administration protocol resulted in elevated plasma levels of BUN and NGAL, indicating kidney dysfunction and injury (Figure 7A-B). Histological analyses revealed significant tubular injury and interstitial fibrosis in the kidneys of cisplatin-treated mice (Figure 7C-F), validating the utility of the three-cycle cisplatin administration protocol as a model for cisplatin-induced nephrotoxicity. We observed that macrophage ablation by PLX3397 provided significant protection against cisplatin-induced kidney dysfunction and prevented both tubular injury and interstitial fibrosis (Figure 7). A pathogenic role of macrophages has been described in various models of kidney disease through studies employing macrophage depletion. For example, in the genetic ablation model using CD11b-DTR mice or treatment with liposomal clodronate, macrophage depletion resulted in improved renal function and mitigated pathological damage in response to acute ischemic kidney injury (*125, 126*) and unilateral ureteral obstruction (*127*). In addition, depletion of kidney resident macrophages attenuated cisplatin-induced kidney fibrosis in mice that were treated with 9mg/kg cisplatin weekly and euthanized three days after the last dose (*59, 124*). In our study, we demonstrated the pathogenic role of macrophages in our clinically relevant mouse model of cisplatin-induced kidney injury and further elucidated the underlying mechanism by which macrophage ablation confers protection against cisplatin-induced nephrotoxicity.

Increased levels of platinum were observed in the kidneys of cisplatin/vehicle-treated mice, while the kidneys of cisplatin/PLX3397-treated mice exhibited significantly less platinum accumulation (Figure 8C). These data suggest that macrophage ablation restricted accumulation of cisplatin in the kidney, thereby offering renal protection by minimizing cisplatin exposure. One possible mechanism for this protection is that macrophages may regulate the expression or function of transporter proteins that mediate uptake or efflux of cisplatin in proximal tubular cells. Renal epithelial cells take up cisplatin through specific transporters, including Cation Transporter 2 (OCT2) and Multidrug and Toxin Extrusion 1 (MATE1). OCT2 is predominantly expressed in the basolateral membrane of proximal tubular cells and mediates cellular uptake of cisplatin (*128, 129*), while MATE1 is expressed in the apical brush-border membrane of renal proximal tubular cells and facilitates cisplatin efflux (*130*). It is plausible that macrophages may regulate expression of these cisplatin transporters in proximal tubular cells in a manner that promotes cisplatin entry. In this model, depletion of macrophages would result in reduced expression of OCT2 (restricting entry of cisplatin into the tubular cells) and/or increased expression of MATE1 (enhancing cisplatin efflux) by the tubular cells, ultimately leading to reduced cisplatin accumulation in the kidney.

### Impact of cisplatin and PLX3397 on resident macrophage populations

Tissue-resident macrophages and microglia rely heavily on CSF1R signaling for their survival (*27, 61, 62*). Two CSF1R inhibitors, PLX3397 and PLX5622, were developed by Plexxicon Inc. (San Francisco, CA). PLX3397 and PLX5622, were initially thought to have minimal impact on peripheral immune cells. However, recent research has demonstrated that both compounds significantly affect tissue-resident macrophages in peripheral organs, such as the peritoneum, lung, and liver (*131–133*). However, to date, there have been no reports investigating the impact of CSF1R inhibitors on resident macrophages in the kidney. We observed that sustained macrophage ablation by PLX3397 resulted in a significant reduction in the number of CX3CR1^GFP^-positive macrophages in the kidney (Figure 8A-B). In contrast to its effect on peripheral organs (e.g. peritoneum, lung, liver, and kidney), PLX3397 had minimal impacts on the populations of leukocytes in the spleen, as assessed by flow cytometry (Supplemental Figure 4-2), consistent with previous reports (*27, 68, 131, 134*).

In contrast to the minimal impact PLX3397 had on immune cell populations in the spleen, treatment with cisplatin did result in a significant reduction in the number of immune cells within the spleen (Supplemental Figure 4-2) and in the cochlea (Figure 1C, Figure 4C). This is consistent with the well-established myelosuppressive effect of cisplatin, which is characterized by reduced bone marrow activity leading to decreased production of red blood cells, white blood cells (immune cells), and platelets. Consequently, cisplatin-induced myelosuppression can lead to hematological toxicities, including anemia, leukopenia, neutropenia, and thrombocytopenia (*6, 135*). Notably, co-treatment of mice with cisplatin and PLX3397 in our study did not result in increased numbers of immune cells in the spleen, suggesting that PLX3397 was not protective against cisplatin-induced myelosuppression (Supplemental Figure 4-2).

In addition to examining the effects of PLX3397 on macrophage numbers, we also examined the effects of cisplatin on numbers of macrophages in the cochlea and kidney. Interestingly, we observed a significant increase in the number of CX3CR1^GFP^-positive cells in the kidney at the end of the cisplatin treatment (Figure 8A-B). This was in contrast to the decrease in CX3CR1^GFP^-positive cells we observed in the cochlea of cisplatin-treated animals (Figure 1C-D, Figure 4C-D). This finding is consistent with previous reports suggesting that peripheral immune cells readily infiltrate the kidney following cisplatin treatment (*83, 122, 124*). Therefore, this difference between the cochlea and kidney may be attributable to peripheral immune cell infiltration into the kidney. Although CX3CR1 is more commonly expressed in macrophages, CX3CR1 expression has also been observed in circulating monocytes, dendritic cells, natural killer cells, and T cells (*59, 136, 137*). Therefore, the observed increase in the CX3CR1^GFP^-positive cells within the kidney during cisplatin treatment may indicate infiltration of a heterogenous population of peripheral immune cells (including macrophages) expressing CX3CR1.

## Conclusion

Our data demonstrate substantial depletion (>88%) of CX3CR1^GFP^-positive macrophages in both the inner ear and kidney using the CSF1R inhibitor PLX3397. Macrophage ablation resulted in protection against cisplatin-induced ototoxicity and nephrotoxicity. Specifically, macrophage ablation preserved hearing function and prevented the loss of OHCs, IHC synapses, and SGNs. In the kidney, we observed substantial protection against cisplatin-induced tubular injury and fibrosis and preservation of kidney function. Additionally, macrophage ablation resulted in a significant reduction in platinum accumulation in both the inner ear and kidney. These data suggest that macrophage ablation confers protection against cisplatin-induced ototoxicity and nephrotoxicity by limiting accumulation of cisplatin in the inner ear and kidney. Therefore, our results are consistent with a model in which macrophages play a role in facilitating cisplatin-induced cytotoxicity by promoting the entry of cisplatin into these organs. Since PLX3397 is FDA-approved to treat tenosynovial giant cell tumors, our data suggest a potential therapeutic approach utilizing PLX3397 to prevent the two most commonly occurring toxicities in cancer patients undergoing cisplatin treatment. A prospective clinical study of PLX3397 that aims to reduce cisplatin-associated ototoxicity and nephrotoxicity in patients with cancer is warranted.

## CRediT authorship contribution statement

**Cathy Yea Won Sung**: Conceptualization, Methodology, Validation, Formal analysis, Investigation, Data Curation, Visualization, Writing – Original Draft, Writing – Review & Editing. **Naoki Hayase**: Formal analysis, Investigation, Data Curation, Visualization, Writing – Review & Editing. **Peter S.T. Yuen**: Supervision. **John Lee**: Investigation, Data Curation, Writing – Review & Editing. **Katharine Fernandez**: Methodology, Investigation, Writing – Review & Editing. **Xuzhen Hu**: Investigation. **Hui Cheng**: Formal analysis, Writing – Review & Editing. **Robert A. Star**: Supervision, Writing – Review & Editing, Project administration, Funding acquisition. **Mark E. Warchol**: Conceptualization, Methodology, Writing – Review & Editing, Funding acquisition. **Lisa L. Cunningham**: Conceptualization, Methodology, Writing – Review & Editing, Supervision, Project administration, Funding acquisition.

## Acknowledgements

This work is supported by NIDCD Division of Intramural Research grant (project number 1 ZIA DC000079), NIDDK Division of Intramural Research grant (project number 1ZIADK043403), and NIDCD R01DC006283. The Authors are grateful to the animal care facility staff of the John Edward Porter Research Neuroscience Center for their ongoing support and exceptional animal care; Tracy Fitzgerald of the NIDCD mouse auditory testing core (project number ZIC DC000080) for training in audiometric techniques; Christopher Silvin of the NCI (previously NIDCD) flow cytometry core and Clint Allen (NCI) for training in the use of flow cytometry and guidance on experimental design; Ukpong Eyo (University of Virginia) for assisting with the PLX3397 pilot experiment, which aimed to determine the ability of PLX3397 to ablate cochlear resident macrophages; Steve Eyles, Director of Mass Spectrometry Core at University of Massachusetts Amherst, for performing ICP-MS analyses. We thank Michael Hoa and Clint Allen for their helpful comments on the manuscript, and Ethan Tyler and Alan Hoofring of the NIH Medical Arts Branch for the illustration in Figure 9.

---

**Supplemental Figure 1.**
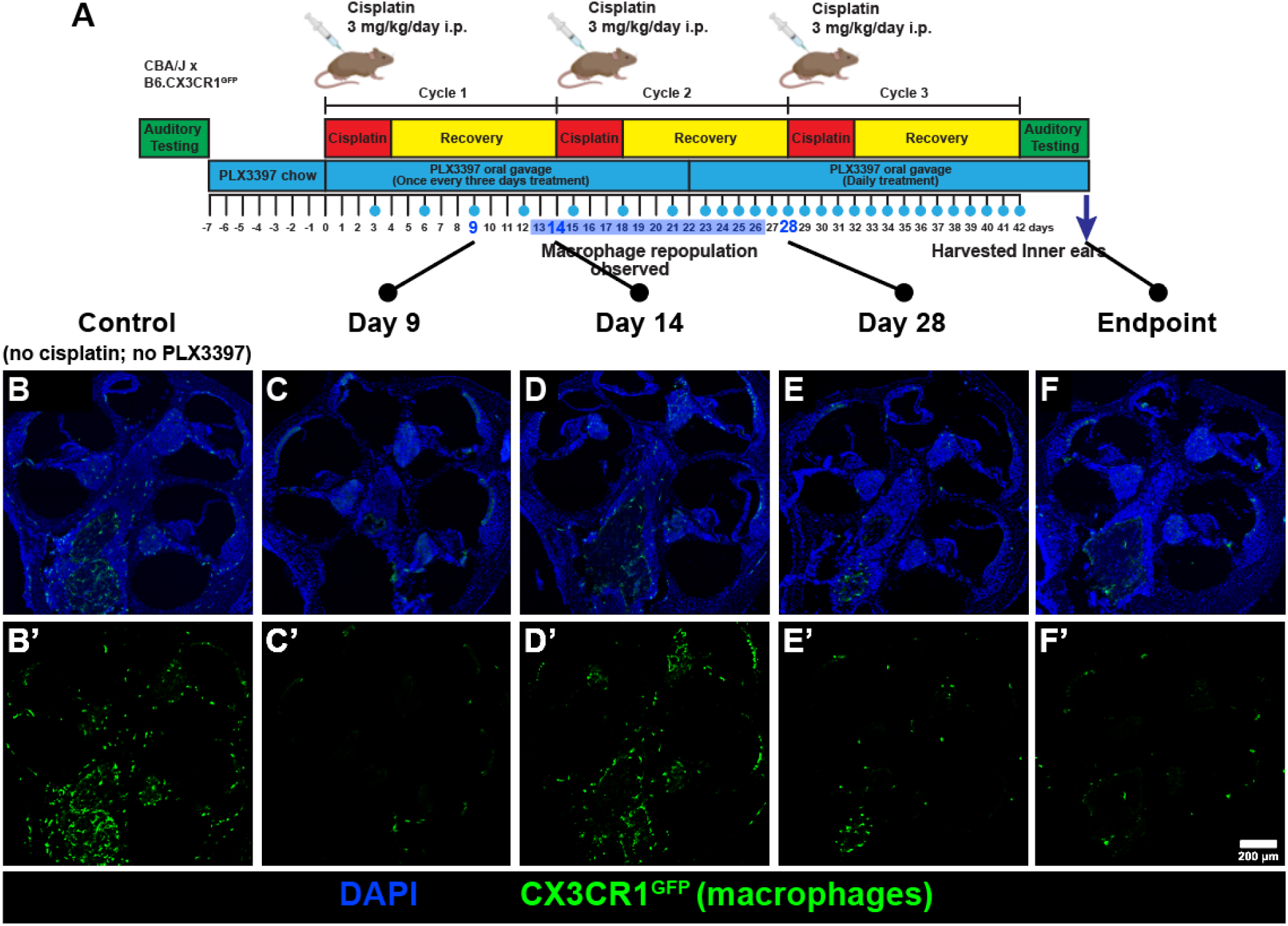
PLX3397 treatment once every three days resulted in partial macrophage repopulation (Experiment 1). (A) Following baseline auditory testing, mice received either control chow or PLX3397-formulated chow for seven days. Subsequently, PLX3397 was administered via oral gavage once every three days, followed by daily dosing initiated after the midpoint of cycle 2 and continued until mice were euthanized after the endpoint auditory testing. (B-F) Cochleae from intermediate time points were harvested to visualize CX3CR1^GFP^-positive macrophages. Nuclei were stained with Hoechst 33342 (blue). (B, B’) Control cochleae were not exposed to cisplatin or PLX3397. (C,C’) Macrophage ablation was efficient at Day 9. (D,D’) Macrophage repopulation was observed at Day 14. (E, E’; F, F’) To address this, the frequency of PLX3397 administration was increased to daily, resulting in subsequent macrophage ablation by day 28. The days on which mice received PLX3397 oral gavage treatment are marked by blue circles in the experimental timeline. *Scale bar, 200 μm*.

**Supplemental Figure 2.**
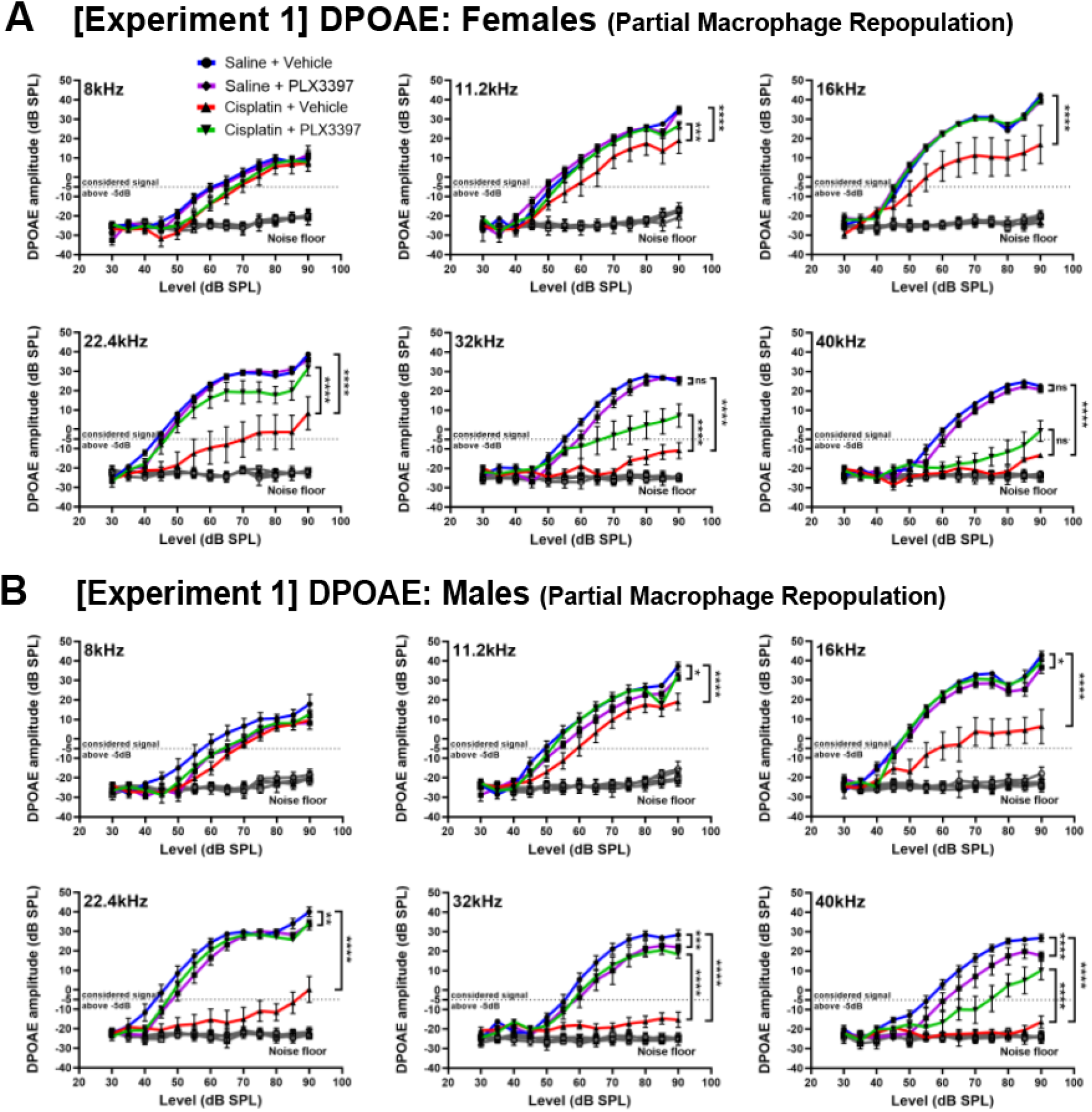
Macrophage ablation resulted in protection against OHC dysfunction in mice of both sexes (Experiment 1). DPOAEs were recorded to assess OHC function in female and male mice. An emission at 2f_1_-f_2_ was considered present when its amplitude exceeded −5dB (dotted line). Cisplatin (red) resulted in loss of DPOAE amplitudes across frequencies. Macrophage ablation using PLX3397 protected against cisplatin-induced loss of DPOAE amplitudes in both (A) female and (B) male mice. Males exhibited a significantly higher level of protection than females at a significance level of 0.1 following PLX3397 treatment at 32kHz and 40kHz. *Grey line indicates the biological noise floor. Statistical analyses were performed using two way-ANOVA with Tukey’s multiple comparisons test (main column effect). Mean±SEM, n=10-16 mice per experimental group. Statistical analyses to identify the difference in protection between males and females were performed using Mixed Effect Modeling with the R package “lmerTest” as described in the Methods*.

**Supplemental Figure 3.**
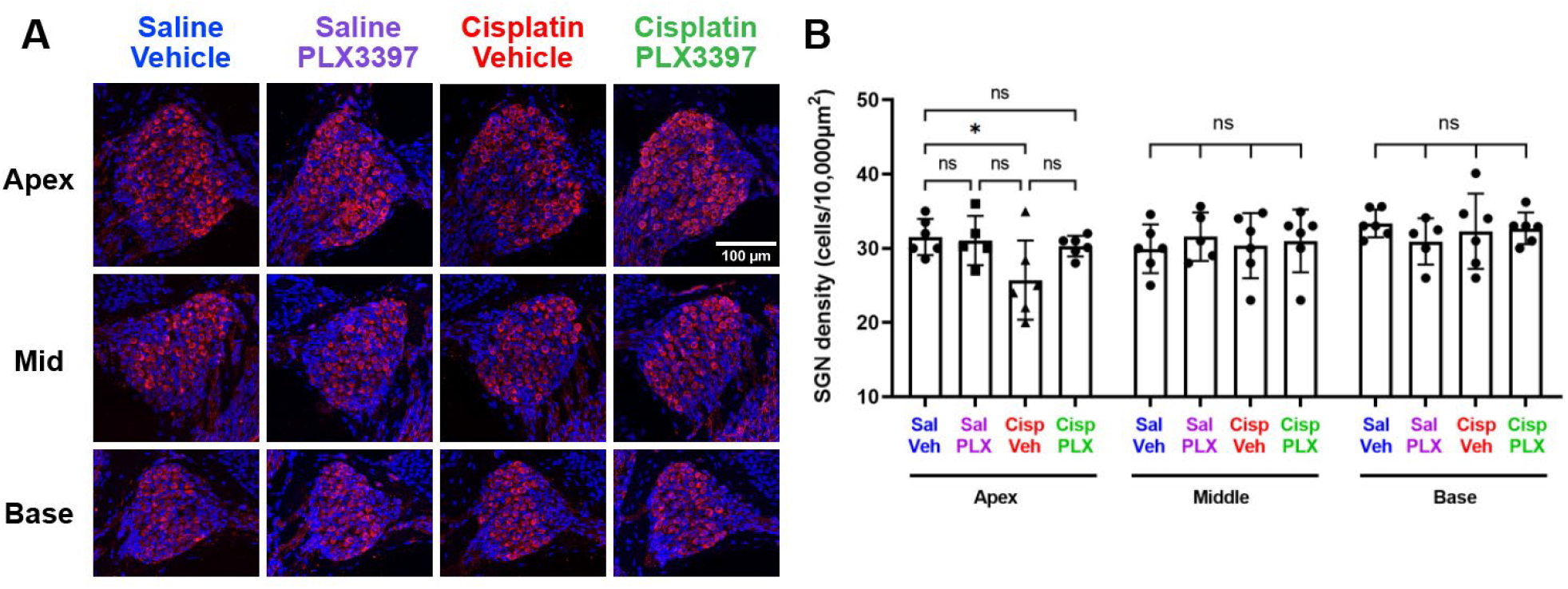
Partial macrophage depletion using PLX3397 protects against cisplatin-induced SGN loss (Experiment 1). Inner ear tissues were harvested following endpoint auditory testing and stained for Tuj-1 (red) to visualize and quantify SGNs. Nuclei were stained with Hoechst 33342 (blue). (A) Representative images and (B) quantification demonstrate that cisplatin-induced SGN loss occurred primarily in the apex, with no significant loss in the middle or basal regions. PLX3397 protected against cisplatin-induced loss of SGNs. (A) *Scale bar, 100 μm. (B) Mean±SD, n=6 inner ears per experimental group. Statistical analysis was performed using one way-ANOVA with Tukey’s multiple comparisons test*.

**Supplemental Figure 4-1.**
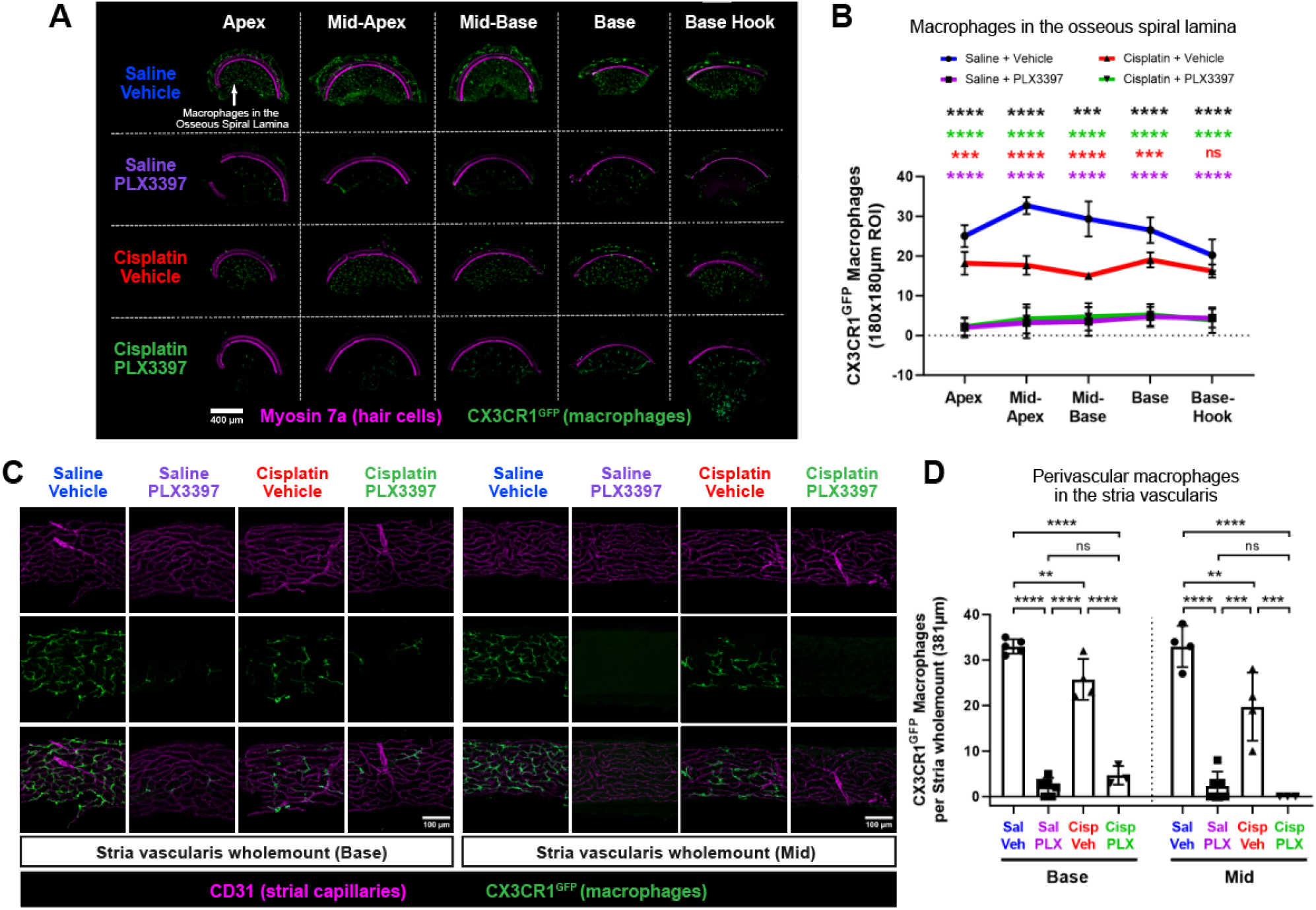
Sustained macrophage ablation using PLX3397 ablates macrophages in the osseous spiral lamina and the stria vascularis (Experiment 2). (A) Representative images and (B) quantitative analysis indicate that daily administration of PLX3397 in Experiment 2 resulted in ablation of macrophages in the osseous spiral lamina overall with the greatest ablation efficiency (92%) in the apex and 80.5% ablation in the basal region of the cochlea, when compared to mice treated with saline/vehicle. In addition, quantitative analysis indicated a significant reduction in macrophages following cisplatin treatment (cisplatin/vehicle-treated mice) compared to mice treated with saline/vehicle, with the middle regions exhibiting the greatest loss (Apex: 27.3%; Mid-Apex: 45.9%; Mid-Base: 48.9%; Base: 28.3%; Base-Hook: 18.9% reduction). *Scale bar, 400 μm*. *Statistical analysis was performed using one-way ANOVA with Tukey’s multiple comparisons test. Statistical comparisons (asterisks or n.s.) are color-coded as described in Methods*. (C-D) Stria vascularis wholemounts were dissected from basal and middle regions of the cochlear lateral wall. (C) Representative images and (D) quantitative analysis indicate that daily administration of PLX3397 in Experiment 2 resulted in ablation of more than 88% of PVMs in the stria vascularis. Cisplatin administration also led to reduction of 25-40% of PVMs compared to the control group (saline/vehicle-treated mice). *Scale bar, 100 μm. (B,D) Mean±SD, n=3-6 cochleae per experimental group. Statistical analysis was performed using one-way ANOVA with Tukey’s multiple comparisons test*.

**Supplemental Figure 4-2.**
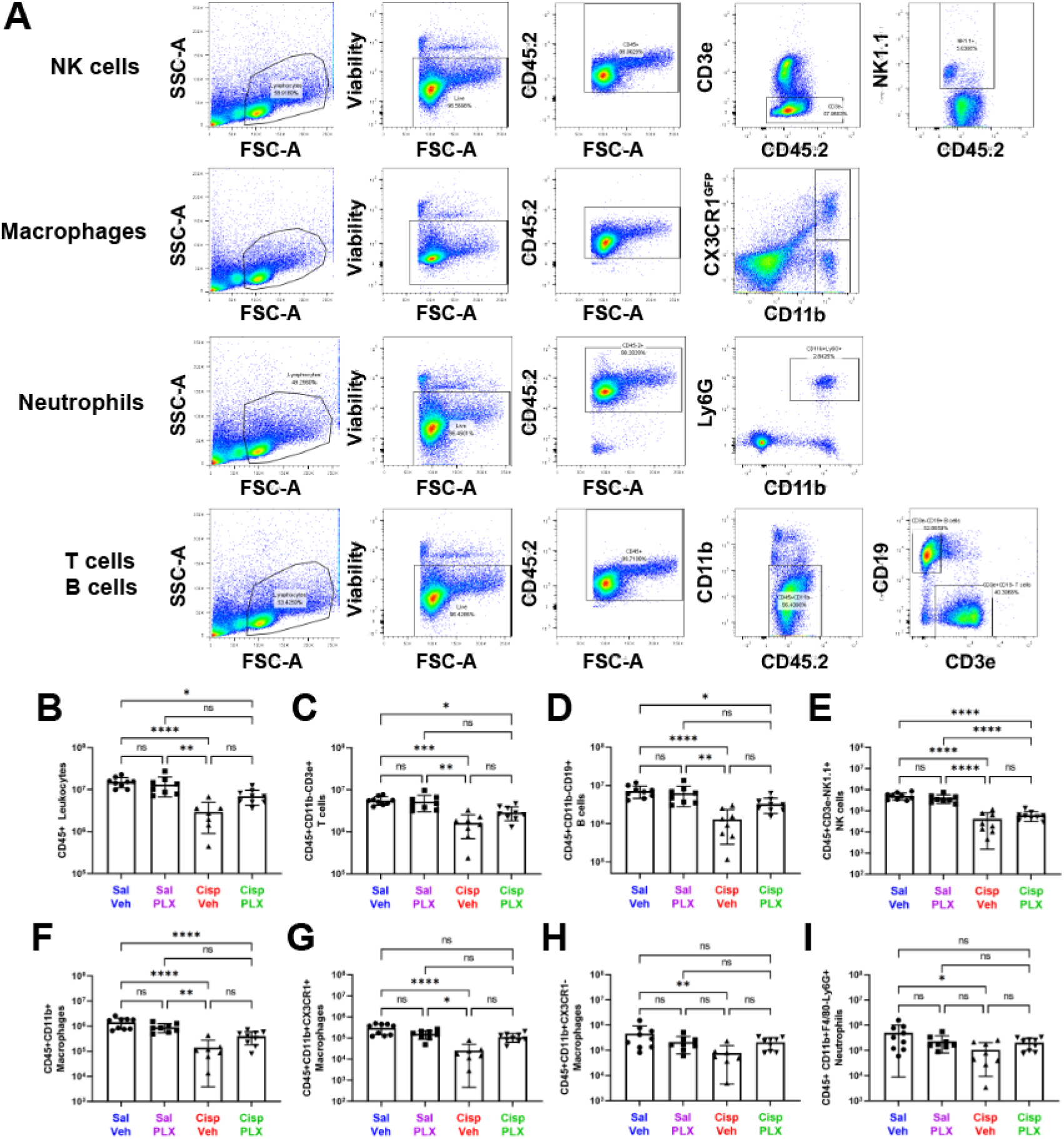
CSF1R inhibition via PLX3397 treatment does not affect peripheral immune cells in the spleen (Experiment 2). Spleens were isolated from CBAJ/CX3CR1^GFP/+^ mice, and single cell suspensions were used for flow cytometric analyses. (A) A representative gating strategy was used to analyze immune cell profiling in the spleen. The absolute numbers of immune cells in the spleen were analyzed: (B) CD45+ Leukocytes, (C) T cells, (D) B cells, (E) NK cells, (F) CD11b+ macrophages, (G) CD11b+ CX3CR1+ macrophages, (H) CD11b+ CX3CR1-macrophages, and (I) neutrophils. Cisplatin reduced the numbers of all analyzed leukocytes. Sustained macrophage ablation via PLX3397 administration did not significantly alter the numbers of leukocytes in the spleen. *Mean±SD, n=7-9 spleens per experimental group. Statistical analysis was performed using Kruskal-Wallis test with Dunn’s multiple comparisons test*.

**Supplemental Figure 5.**
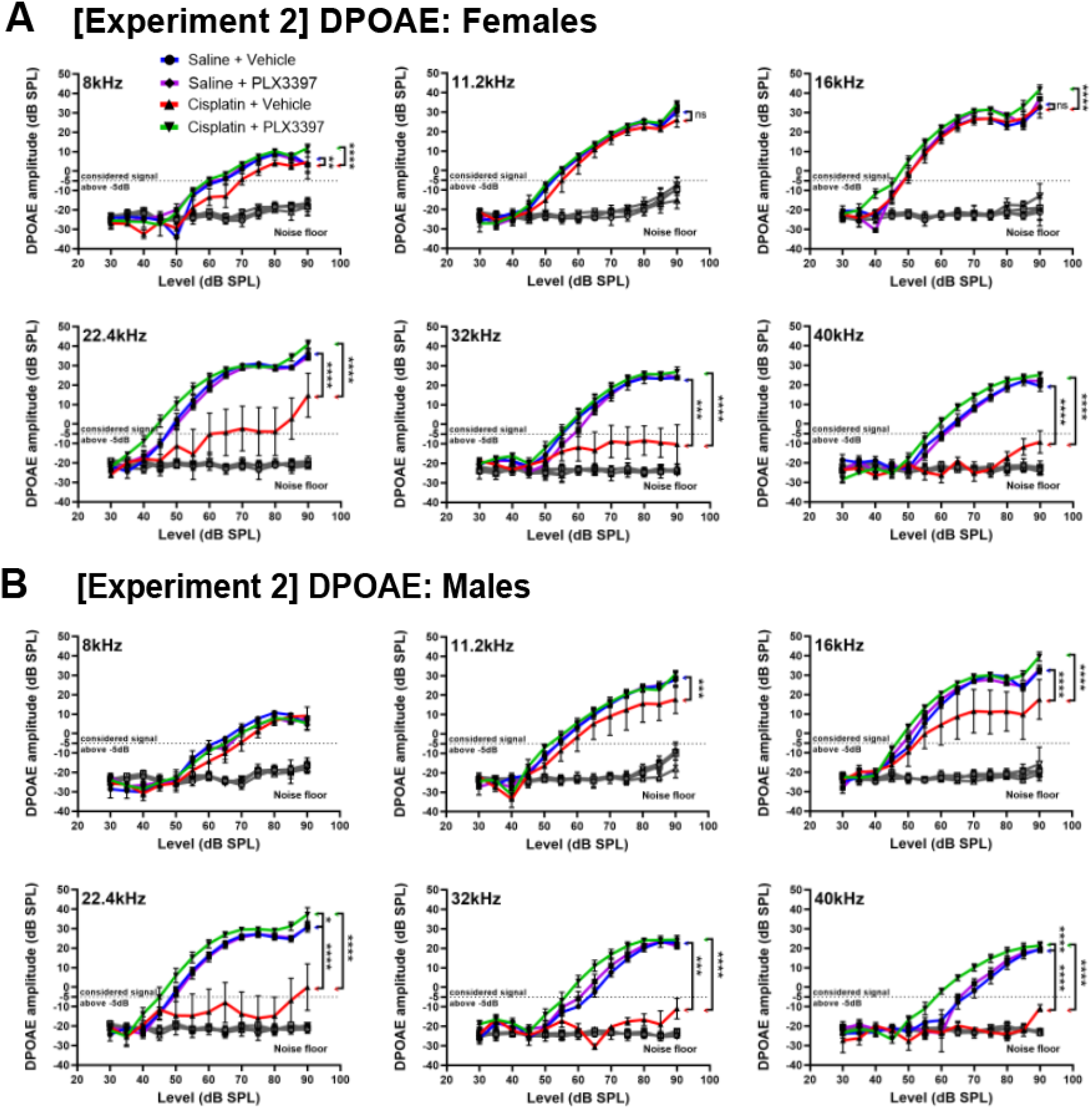
Sustained macrophage ablation using PLX3397 resulted in complete protection against cisplatin-induced OHC dysfunction in both male and female mice (Experiment 2). OHC function was evaluated by DPOAE in (A) female and (B) male mice. In cisplatin/vehicle-treated mice, female mice exhibited a significant reduction in DPOAE amplitudes at frequencies of 22.4 kHz and above, while male mice showed reductions at 11.2kHz and above. PLX3397 resulted in complete protection against cisplatin-induced OHC dysfunction in both female and male mice. *DPOAEs were considered present at 2f_1_-f_2_ when the DPOAE amplitude surpassed the −5dB threshold (dotted line). The grey line represents the biological noise floor. Data are shown as Mean±SEM, n=8-9 mice per experimental group. Statistical analysis was performed using two way-ANOVA with Tukey’s multiple comparisons test (main column effect)*.

**Supplemental Figure 6.**
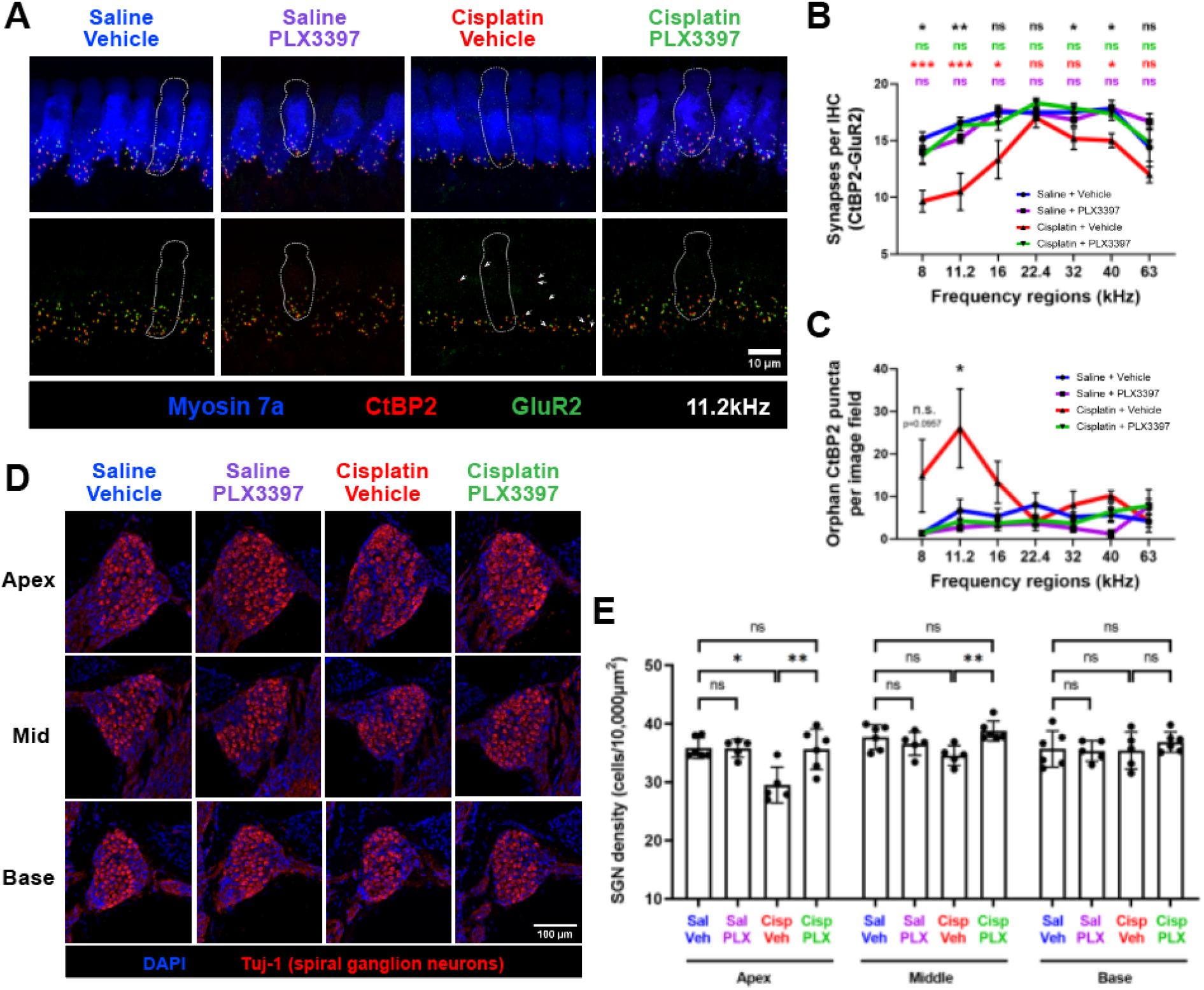
Sustained macrophage ablation protects against cisplatin-induced loss of synapses and SGNs (Experiment 2). (A-C) Cochlear wholemounts were stained for myosin 7a (blue), CtBP2 (pre-synaptic ribbon), and GluR2 (post-synaptic glutamate receptor). (A) Representative images and quantitative analysis reveal a significant loss of (B) CtBP2-GluR2 juxtaposed synapses at the apical-to-mid cochlear regions (8, 11.2, 16kHz) following cisplatin treatment. (A) Orphan CtBP2 pre-synaptic punctae were observed in cisplatin/vehicle-treated mice (white arrows) and (C) quantified. Mice co-administered with cisplatin and PLX3397 exhibited complete preservation of synapses. *(A) Scale bar, 10 μm. (B-C) Data are shown as Mean±SEM, n=6 cochleae per experimental group. P values were calculated using one-way ANOVA with Tukey’s multiple comparisons test. Statistical comparisons (asterisks or n.s.) are color-coded as outlined in Methods.* (D-E) Mid-modiolar cochlear sections were stained for Tuj-1 to visualize and quantify SGNs. Nuclei were stained with Hoechst 33342 (blue). (D) Representative images and (E) quantitative analysis demonstrate cisplatin-induced SGN loss in the apical and middle regions of the cochlea. PLX3397 provided complete protection against cisplatin-induced loss of SGNs. *(D) Scale bar, 100 μm. (E) Data are shown as Mean±SD, n=5-6 cochleae per experimental group. P values were calculated using one-way ANOVA with Tukey’s multiple comparisons test*.

**Supplemental Table 1.**
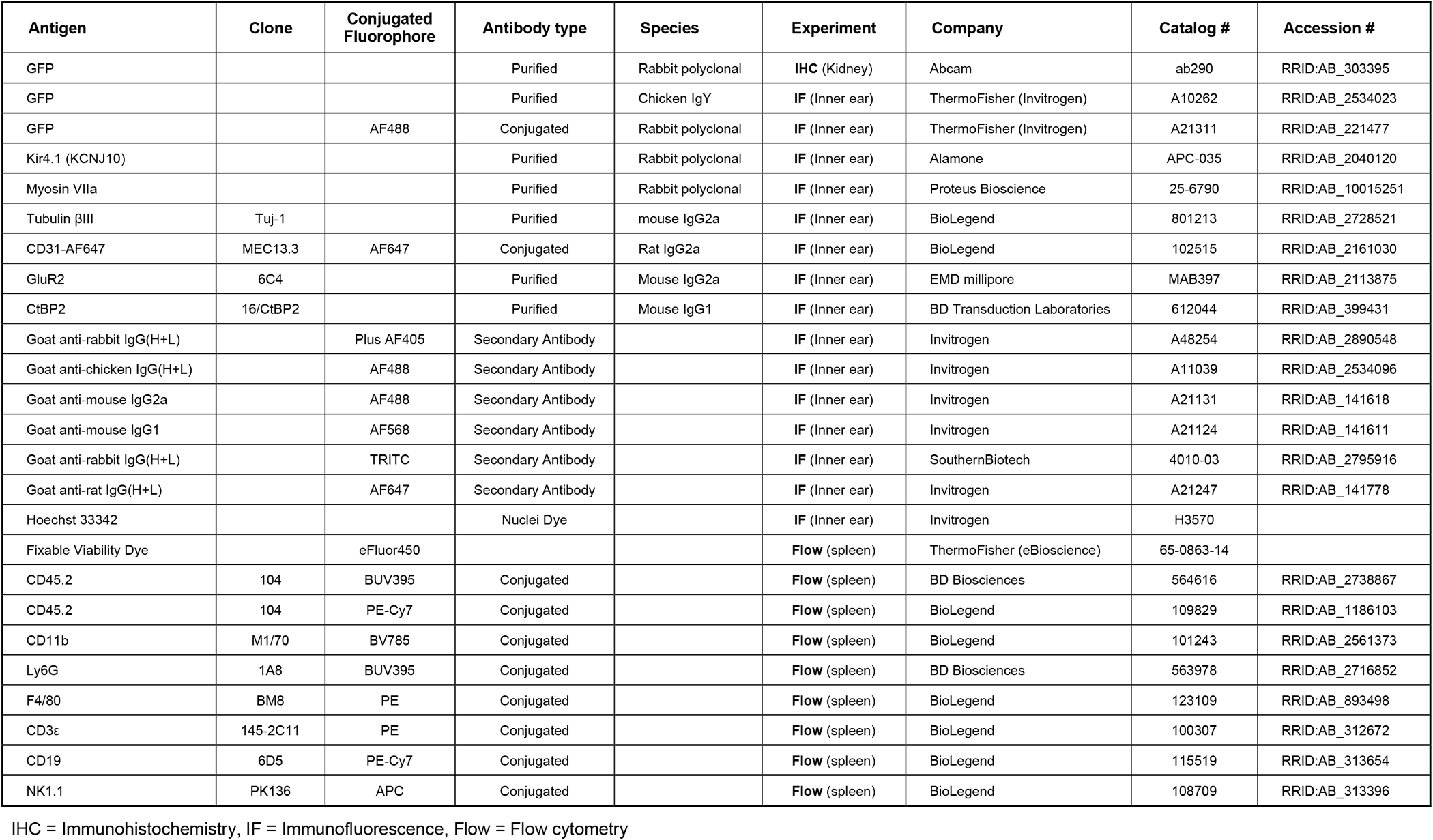
Reagents and antibodies used in this study.

