## Supplemental Figure 1 for "Macrophage Depletion Protects Against Cisplatin-Induced Ototoxicity and Nephrotoxicity"

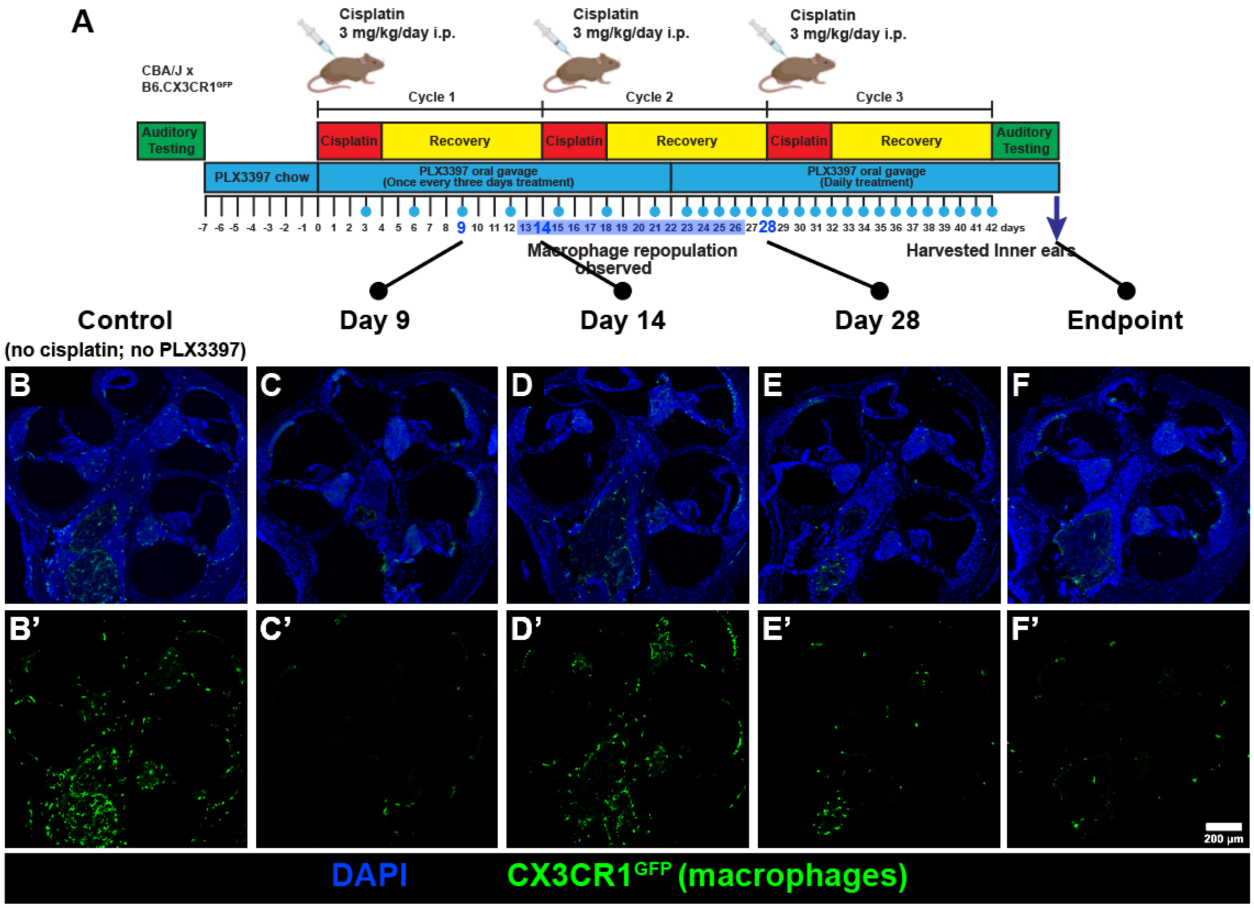


**Supplemental Figure 1. PLX3397 treatment once every three days resulted in partial macrophage repopulation (Experiment 1).** (A) Following baseline auditory testing, mice received either control chow or PLX3397-formulated chow for seven days. Subsequently, PLX3397 was administered via oral gavage once every three days, followed by daily dosing initiated after the midpoint of cycle 2 and continued until mice were euthanized after the endpoint auditory testing. (B-F) Cochleae from intermediate time points were harvested to visualize CX3CR1$^{GFP}$-positive macrophages. Nuclei were stained with Hoechst 33342 (blue). (B, B') Control cochleae were not exposed to cisplatin or PLX3397. (C,C') Macrophage ablation was efficient at Day 9. (D,D') Macrophage repopulation was observed at Day 14. (E, E'; F, F') To address this, the frequency of PLX3397 administration was increased to daily, resulting in subsequent macrophage ablation by day 28. The days on which mice received PLX3397 oral gavage treatment are marked by blue circles in the experimental timeline. *Scale bar, 200 µm*.
