## Supplemental Figure 2 for "Macrophage Depletion Protects Against Cisplatin-Induced Ototoxicity and Nephrotoxicity"

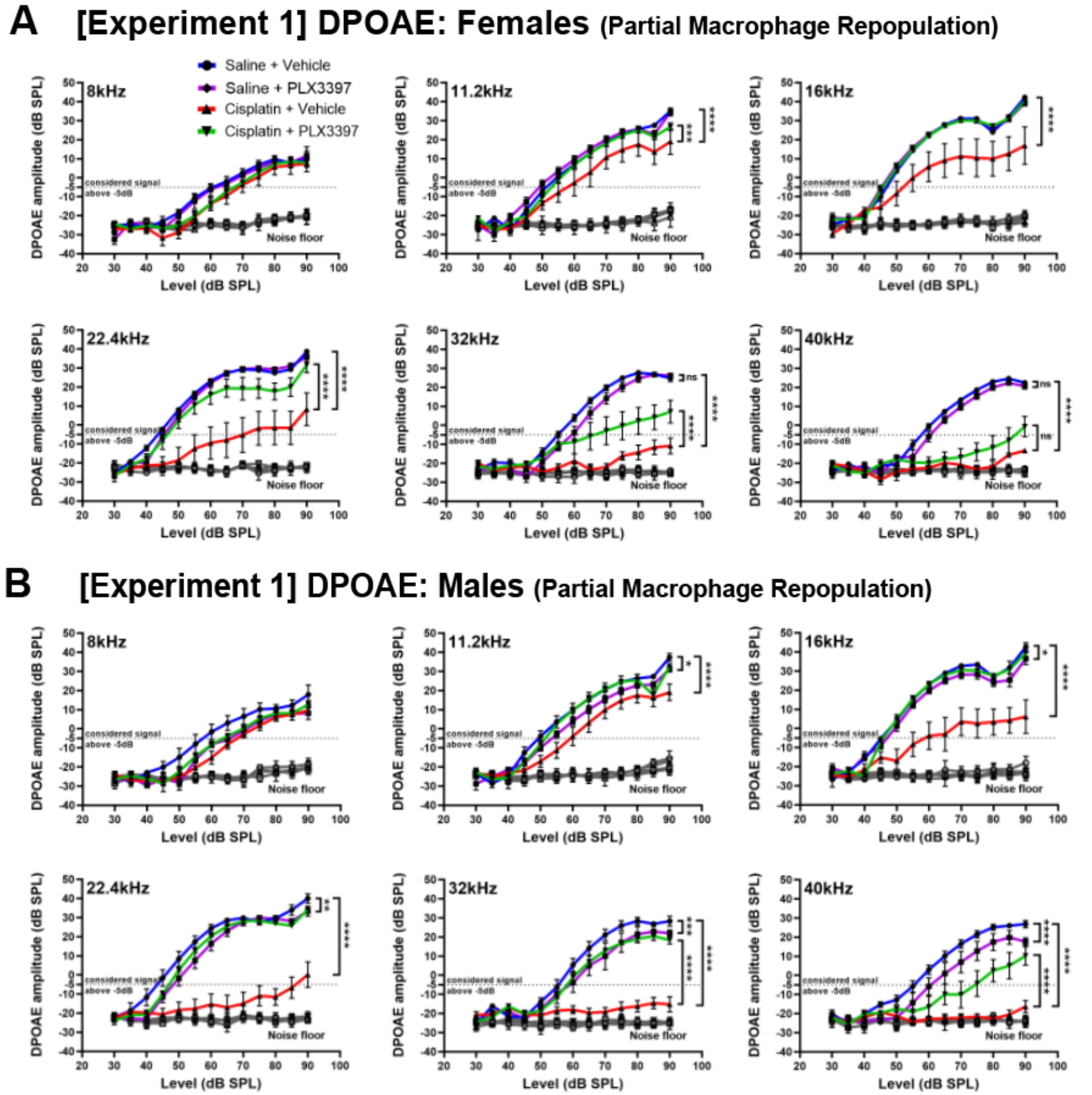


**Supplemental Figure 2. Macrophage ablation resulted in protection against OHC dysfunction in mice of both sexes (Experiment 1).** DPOAEs were recorded to assess OHC function in female and male mice. An emission at $2f_1$-$f_2$ was considered present when its amplitude exceeded -5dB (dotted line). Cisplatin (red) resulted in loss of DPOAE amplitudes across frequencies. Macrophage ablation using PLX3397 protected against cisplatin-induced loss of DPOAE amplitudes in both (A) female and (B) male mice. Males exhibited a significantly higher level of protection than females at a significance level of 0.1 following PLX3397 treatment at 32kHz and 40kHz. *Grey line indicates the biological noise floor. Statistical analyses were performed using two way-ANOVA with Tukey's multiple comparisons test (main column effect). Mean±SEM, n=10-16 mice per experimental group. Statistical analyses to identify the difference in protection between males and females were performed using Mixed Effect Modeling with the R package "lmerTest" as described in the Methods.*
