## Supplemental Figure 3 for "Macrophage Depletion Protects Against Cisplatin-Induced Ototoxicity and Nephrotoxicity"

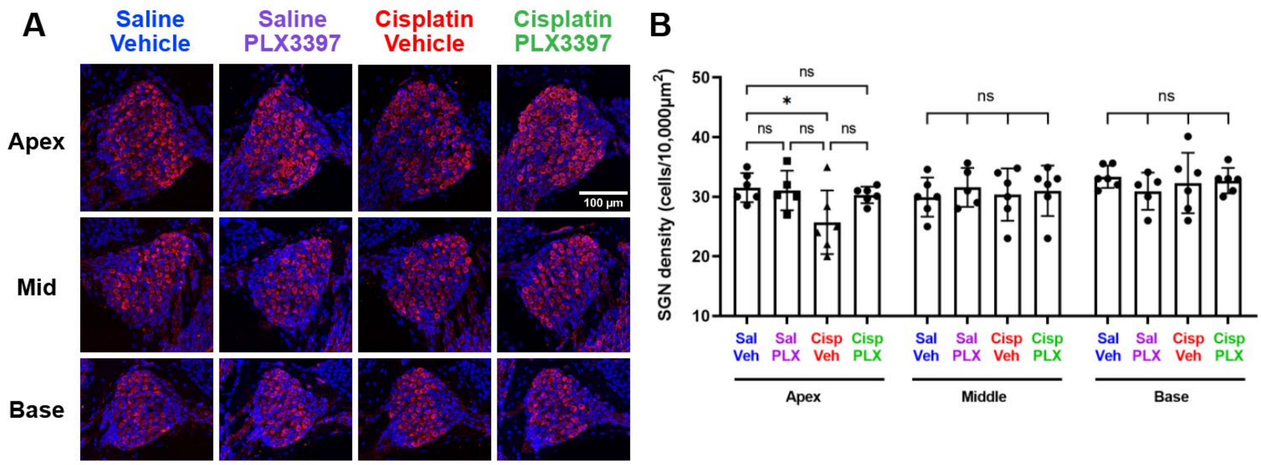


**Supplemental Figure 3. Partial macrophage depletion using PLX3397 protects against cisplatin-induced SGN loss (Experiment 1).** Inner ear tissues were harvested following endpoint auditory testing and stained for Tuj-1 (red) to visualize and quantify SGNs. Nuclei were stained with Hoechst 33342 (blue). (A) Representative images and (B) quantification demonstrate that cisplatin-induced SGN loss occurred primarily in the apex, with no significant loss in the middle or basal regions. PLX3397 protected against cisplatin-induced loss of SGNs. (A) *Scale bar, 100 µm. (B) Mean±SD, n=6 inner ears per experimental group. Statistical analysis was performed using one way-ANOVA with Tukey's multiple comparisons test.*
