## Supplemental Figure 4-1 for "Macrophage Depletion Protects Against Cisplatin-Induced Ototoxicity and Nephrotoxicity"

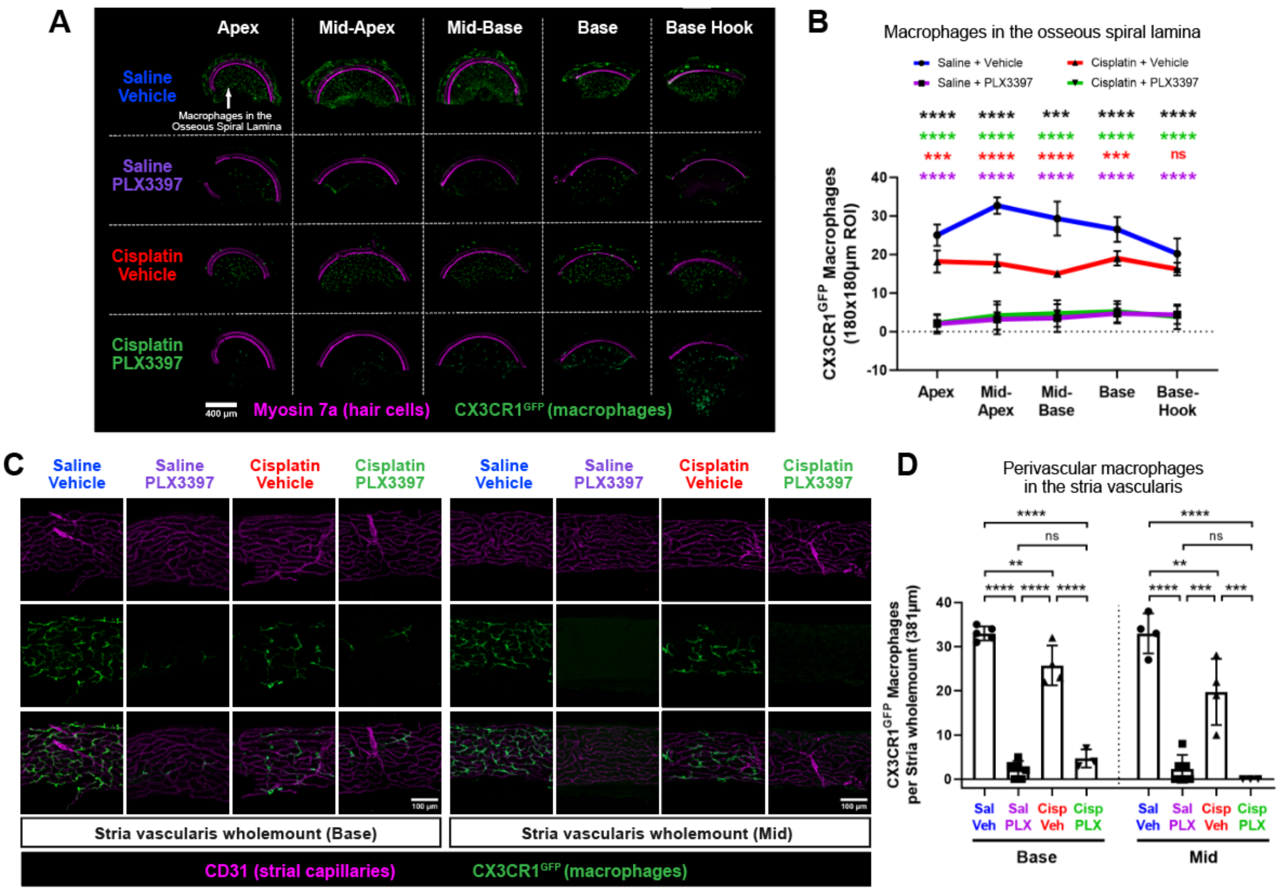


**Supplemental Figure 4-1. Sustained macrophage ablation using PLX3397 ablates macrophages in the osseous spiral lamina and the stria vascularis (Experiment 2).** (A) Representative images and (B) quantitative analysis indicate that daily administration of PLX3397 in Experiment 2 resulted in ablation of macrophages in the osseous spiral lamina overall with the greatest ablation efficiency (92%) in the apex and 80.5% ablation in the basal region of the cochlea, when compared to mice treated with saline/vehicle. In addition, quantitative analysis indicated a significant reduction in macrophages following cisplatin treatment (cisplatin/vehicle-treated mice) compared to mice treated with saline/vehicle, with the middle regions exhibiting the greatest loss (Apex: 27.3%; Mid-Apex: 45.9%; Mid-Base: 48.9%; Base: 28.3%; Base-Hook: 18.9% reduction). *Scale bar, 400 µm. Statistical analysis was performed using one-way ANOVA with Tukey's multiple comparisons test. Statistical comparisons (asterisks or n.s.) are color-coded as described in Methods.* (C-D) Stria vascularis wholemounts were dissected from basal and middle regions of the cochlear lateral wall. (C) Representative images and (D) quantitative analysis indicate that daily administration of PLX3397 in Experiment 2 resulted in ablation of more than 88% of PVMs in the stria vascularis. Cisplatin administration also led to reduction of 25-40% of PVMs compared to the control group (saline/vehicle-treated mice). *Scale bar, 100 µm. (B,D) Mean±SD, n=3-6 cochleae per experimental group. Statistical analysis was performed using one-way ANOVA with Tukey's multiple comparisons test.*
