## Supplemental Figure 4-2 for "Macrophage Depletion Protects Against Cisplatin-Induced Ototoxicity and Nephrotoxicity"

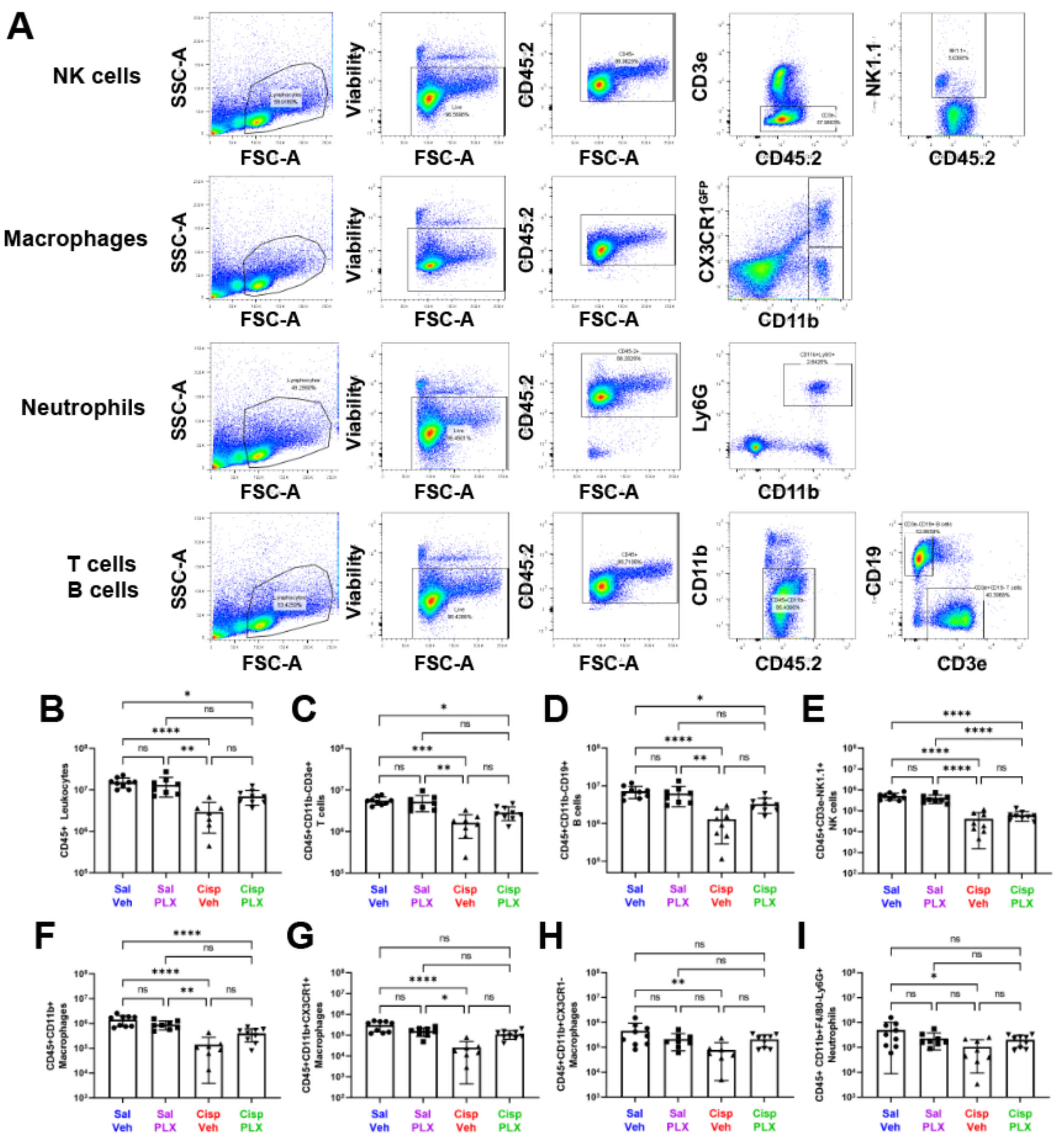


**Supplemental Figure 4-2. CSF1R inhibition via PLX3397 treatment does not affect peripheral immune cells in the spleen (Experiment 2).** Spleens were isolated from CBAJ/CX3CR1$^{GFP/+}$ mice, and single cell suspensions were used for flow cytometric analyses. (A) A representative gating strategy was used to analyze immune cell profiling in the spleen. The absolute numbers of immune cells in the spleen were analyzed: (B) CD45+ Leukocytes, (C) T cells, (D) B cells, (E) NK cells, (F) CD11b+ macrophages, (G) CD11b+ CX3CR1+ macrophages, (H) CD11b+ CX3CR1- macrophages, and (I) neutrophils. Cisplatin reduced the numbers of all analyzed leukocytes. Sustained macrophage ablation via PLX3397 administration did not significantly alter the numbers of leukocytes in the spleen. *Mean±SD, n=7-9 spleens per experimental group. Statistical analysis was performed using Kruskal-Wallis test with Dunn's multiple comparisons test.*
