## Supplemental Figure 5 for "Macrophage Depletion Protects Against Cisplatin-Induced Ototoxicity and Nephrotoxicity"

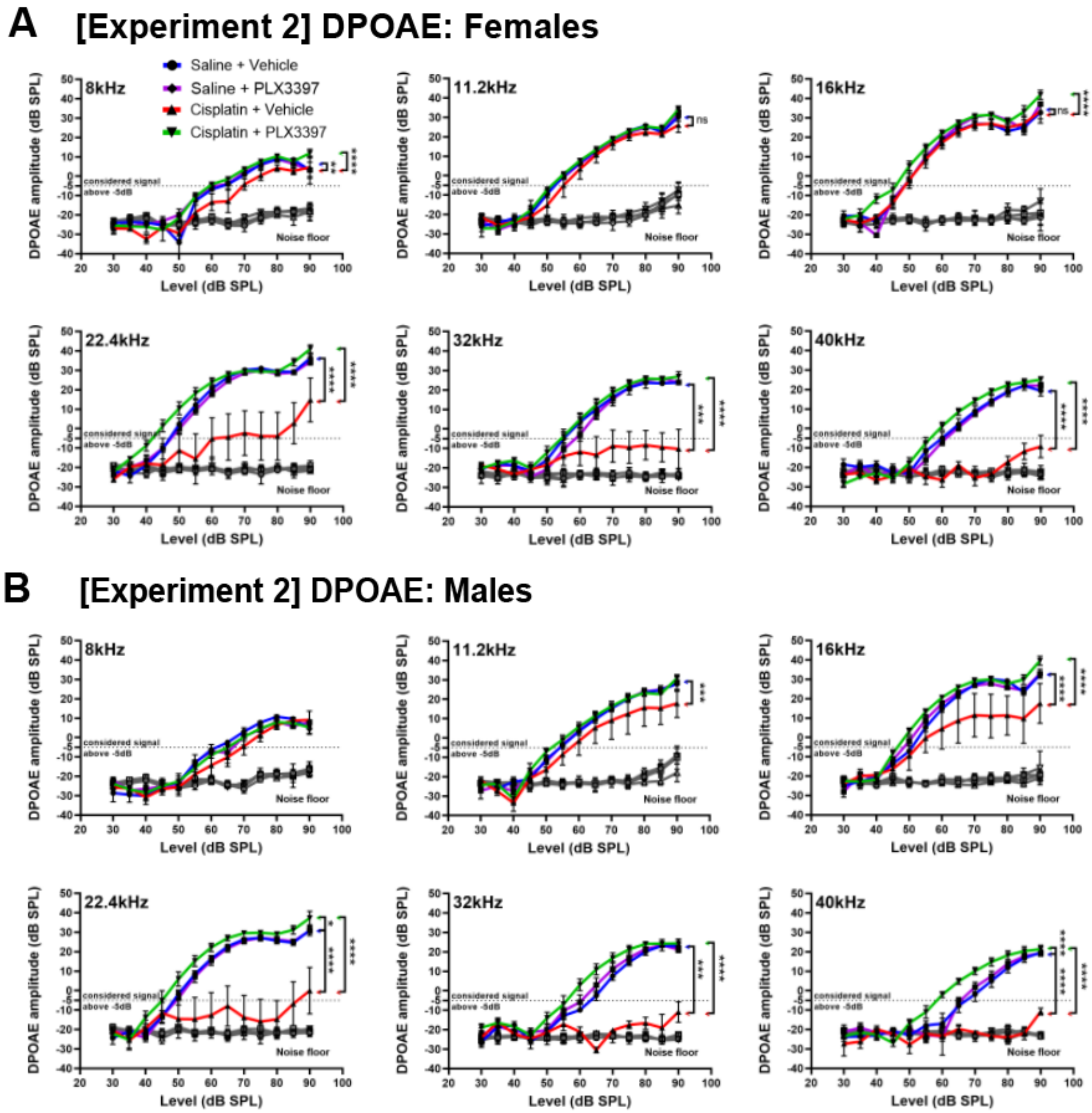


**Supplemental Figure 5. Sustained macrophage ablation using PLX3397 resulted in complete protection against cisplatin-induced OHC dysfunction in both male and female mice (Experiment 2).** OHC function was evaluated by DPOAE in (A) female and (B) male mice. In cisplatin/vehicle-treated mice, female mice exhibited a significant reduction in DPOAE amplitudes at frequencies of 22.4 kHz and above, while male mice showed reductions at 11.2kHz and above. PLX3397 resulted in complete protection against cisplatin-induced OHC dysfunction in both female and male mice. *DPOAEs were considered present at $2f_1$-$f_2$ when the DPOAE amplitude surpassed the -5dB threshold (dotted line). The grey line represents the biological noise floor. Data are shown as Mean±SEM, n=8-9 mice per experimental group. Statistical analysis was performed using two way-ANOVA with Tukey's multiple comparisons test (main column effect).*
