## Supplemental Figure 6 for "Macrophage Depletion Protects Against Cisplatin-Induced Ototoxicity and Nephrotoxicity"

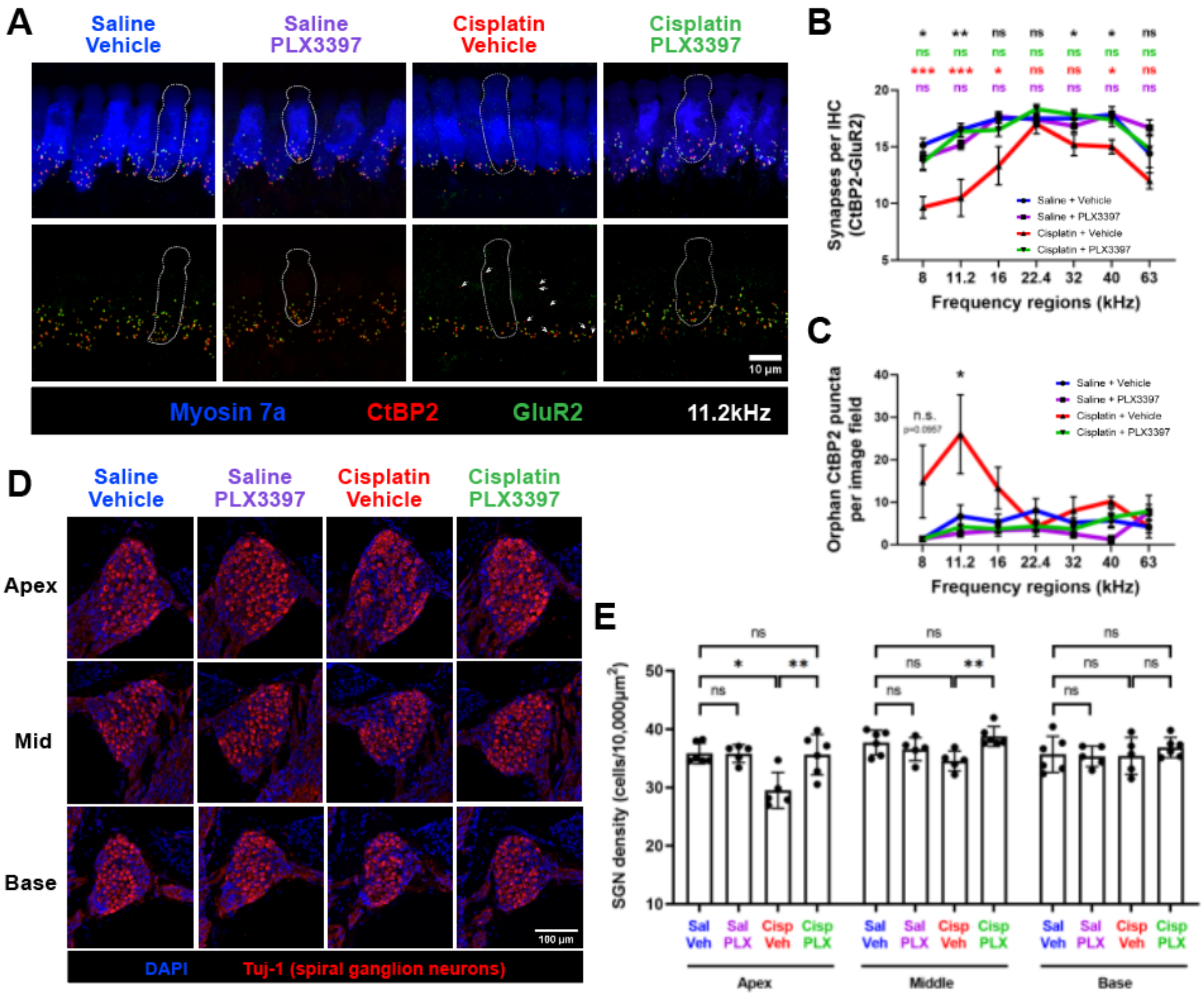


**Supplemental Figure 6. Sustained macrophage ablation protects against cisplatin-induced loss of synapses and SGNs (Experiment 2).** (A-C) Cochlear wholemounts were stained for myosin 7a (blue), CtBP2 (pre-synaptic ribbon), and GluR2 (post-synaptic glutamate receptor). (A) Representative images and quantitative analysis reveal a significant loss of (B) CtBP2-GluR2 juxtaposed synapses at the apical-to-mid cochlear regions (8, 11.2, 16kHz) following cisplatin treatment. (A) Orphan CtBP2 pre-synaptic punctae were observed in cisplatin/vehicle-treated mice (white arrows) and (C) quantified. Mice co-administered with cisplatin and PLX3397 exhibited complete preservation of synapses. *(A) Scale bar, 10 µm. (B-C) Data are shown as Mean±SEM, n=6 cochleae per experimental group. P values were calculated using one-way ANOVA with Tukey's multiple comparisons test. Statistical comparisons (asterisks or n.s.) are color-coded as outlined in Methods.* (D-E) Mid-modiolar cochlear sections were stained for Tuj-1 to visualize and quantify SGNs. Nuclei were stained with Hoechst 33342 (blue). (D) Representative images and (E) quantitative analysis demonstrate cisplatin-induced SGN loss in the apical and middle regions of the cochlea. PLX3397 provided complete protection against cisplatin-induced loss of SGNs. *(D) Scale bar, 100 µm. (E) Data are shown as Mean±SD, n=5-6 cochleae per experimental group. P values were calculated using one-way ANOVA with Tukey's multiple comparisons test.*
