## Supplemental Table 1 for "Macrophage Depletion Protects Against Cisplatin-Induced Ototoxicity and Nephrotoxicity"

Supplemental Table 1. Reagents and antibodies used in this study.

| Antigen | Clone | Conjugated Fluorophore | Antibody type | Species | Experiment | Company | Catalog # | Accession # |
|---|---|---|---|---|---|---|---|---|
| GFP | | | Purified | Rabbit polyclonal | **IHC** (Kidney) | Abcam | ab290 | RRID:AB_303395 |
| GFP | | | Purified | Chicken IgY | **IF** (Inner ear) | ThermoFisher (Invitrogen) | A10262 | RRID:AB_2534023 |
| GFP | | AF488 | Conjugated | Rabbit polyclonal | **IF** (Inner ear) | ThermoFisher (Invitrogen) | A21311 | RRID:AB_221477 |
| Kir4.1 (KCNJ10) | | | Purified | Rabbit polyclonal | **IF** (Inner ear) | Alamone | APC-035 | RRID:AB_2040120 |
| Myosin VIIa | | | Purified | Rabbit polyclonal | **IF** (Inner ear) | Proteus Bioscience | 25-6790 | RRID:AB_10015251 |
| Tubulin βIII | Tuj-1 | | Purified | mouse IgG2a | **IF** (Inner ear) | BioLegend | 801213 | RRID:AB_2728521 |
| CD31-AF647 | MEC13.3 | AF647 | Conjugated | Rat IgG2a | **IF** (Inner ear) | BioLegend | 102515 | RRID:AB_2161030 |
| GluR2 | 6C4 | | Purified | Mouse IgG2a | **IF** (Inner ear) | EMD millipore | MAB397 | RRID:AB_2113875 |
| CtBP2 | 16/CtBP2 | | Purified | Mouse IgG1 | **IF** (Inner ear) | BD Transduction Laboratories | 612044 | RRID:AB_399431 |
| Goat anti-rabbit IgG(H+L) | | Plus AF405 | Secondary Antibody | | **IF** (Inner ear) | Invitrogen | A48254 | RRID:AB_2890548 |
| Goat anti-chicken IgG(H+L) | | AF488 | Secondary Antibody | | **IF** (Inner ear) | Invitrogen | A11039 | RRID:AB_2534096 |
| Goat anti-mouse IgG2a | | AF488 | Secondary Antibody | | **IF** (Inner ear) | Invitrogen | A21131 | RRID:AB_141618 |
| Goat anti-mouse IgG1 | | AF568 | Secondary Antibody | | **IF** (Inner ear) | Invitrogen | A21124 | RRID:AB_141611 |
| Goat anti-rabbit IgG(H+L) | | TRITC | Secondary Antibody | | **IF** (Inner ear) | SouthernBiotech | 4010-03 | RRID:AB_2795916 |
| Goat anti-rat IgG(H+L) | | AF647 | Secondary Antibody | | **IF** (Inner ear) | Invitrogen | A21247 | RRID:AB_141778 |
| Hoechst 33342 | | | Nuclei Dye | | **IF** (Inner ear) | Invitrogen | H3570 | |
| Fixable Viability Dye | | eFluor450 | | | **Flow** (spleen) | ThermoFisher (eBioscience) | 65-0863-14 | |
| CD45.2 | 104 | BUV395 | Conjugated | | **Flow** (spleen) | BD Biosciences | 564616 | RRID:AB_2738867 |
| CD45.2 | 104 | PE-Cy7 | Conjugated | | **Flow** (spleen) | BioLegend | 109829 | RRID:AB_1186103 |
| CD11b | M1/70 | BV785 | Conjugated | | **Flow** (spleen) | BioLegend | 101243 | RRID:AB_2561373 |
| Ly6G | 1A8 | BUV395 | Conjugated | | **Flow** (spleen) | BD Biosciences | 563978 | RRID:AB_2716852 |
| F4/80 | BM8 | PE | Conjugated | | **Flow** (spleen) | BioLegend | 123109 | RRID:AB_893498 |
| CD3ε | 145-2C11 | PE | Conjugated | | **Flow** (spleen) | BioLegend | 100307 | RRID:AB_312672 |
| CD19 | 6D5 | PE-Cy7 | Conjugated | | **Flow** (spleen) | BioLegend | 115519 | RRID:AB_313654 |
| NK1.1 | PK136 | APC | Conjugated | | **Flow** (spleen) | BioLegend | 108709 | RRID:AB_313396 |

IHC = Immunohistochemistry, IF = Immunofluorescence, Flow = Flow cytometry
